# Neuronal selectivity for multiple features in the primary visual cortex

**DOI:** 10.1101/2022.07.18.500396

**Authors:** Wenqing Wei, Benjamin Merkt, Stefan Rotter

**Affiliations:** Bernstein Center Freiburg, University of Freiburg, Freiburg, Germany; Faculty of Biology, University of Freiburg, Freiburg, Germany

## Abstract

Neurons in rodent primary visual cortex are simultaneously tuned to several stimulus features, including orientation and spatial frequency of moving gratings used in experiments. Light-induced signals emitted by retinal ganglion cells (RGC) are relayed to the primary visual cortex (V1) via cells in the dorsal lateral geniculate nucleus (dLGN). However, there is currently no agreement on which thalamocortical transformation leads to the neuronal tuning curves observed in experiments. Here, we outline a model that explains the emergence of feature-specific neural responses as the result of a two-step integration process: First, the compound input to cortical neurons comes from a set of retinal sensors randomly placed in the receptive field. Second, the cortical responses to the combined input are shaped by the rectification caused by the spike threshold of the neurons. We performed numerical simulations of a thalamocortical network stimulated by moving gratings and found that simultaneous tuning to orientation and spatial frequency results from this spatio-temporal integration process. We also show how this tuning is related to the complex structure of the receptive fields that reflect the input. We conclude that different types of feature selectivity arise naturally from random thalamocortical projections. Moreover, we describe in detail the underlying neural mechanism.

## Introduction

Most neurons in the primary visual cortex (V1) of mammals respond selectively to the orientation of light bars, edges of objects, and oriented gratings (***Hubel and Wiesel, 1962***; ***Ferster and Miller, 2000***; ***Niell and Stryker, 2008***; ***McLaughlin et al., 2000***). Orientation selectivity (OS) is the result of computations in neural circuits. It has been considered as a prototypical example of such sensory computations since it was first characterized by ***Hubel and Wiesel (1962)***. Although a large number of experimental and theoretical approaches have been suggested, the exact neuronal mechanisms underlying the emergence of OS are still controversial. In some mammalian species, neighboring cortical neurons across all layers have similar orientation preferences (***Hubel and Wiesel, 1962***; ***Kremkow et al., 2016***; ***Hubel and Wiesel, 1977***; ***Blasdel and Salama, 1986***). In other species, where there is no such order, individual V1 neurons still exhibit strong orientation tuning (***Ohki et al., 2005***; ***Niell and Stryker, 2008***; ***Hofer et al., 2011***). Therefore, it is not clear whether the same mechanism for the emergence of OS applies to all species.

In the feedforward model originally proposed by ***Hubel and Wiesel (1962)***, the receptive fields of dorsal lateral geniculate nucleus (dLGN) neurons converging to a single V1 neuron are assumed to be lined up in the visual field. Under certain conditions, this arrangement of inputs implies an elongated receptive field of the V1 target neuron, which then exhibits selectivity for a stimulus of matching orientation. This concept, however, cannot explain the pronounced dependence of orientation tuning on the spatial frequency of the grating used for stimulation (***Ayzenshtat et al., 2016***).

In addition, this model requires a mechanism to establish the specific arrangement of receptive fields during development, possibly driven by visual experience. Interestingly, however, mouse V1 neurons exhibit OS already when they open their eyes for the first time and V1 circuits are not yet fully maturated (***Ko et al., 2013***). Therefore, an alternative mechanism that does not depend on precisely arranged thalamocortical projections might underly the emergence of OS. In fact, several alternative such mechanisms have been suggested in the past (***Priebe, 2016***; ***Jang et al., 2020***).

In previous theoretical work (***Hansel and van Vreeswijk, 2012***; ***Pehlevan and Sompolinsky, 2014***; ***Sadeh et al., 2014***; ***Sadeh and Rotter, 2015***), it was pointed out that highly selective and contrast-invariant neuronal responses robustly emerge in inhibition-dominated random recurrent networks. In these models, it was assumed that each V1 neuron receives a mix of multiple thalamic inputs with a weak bias for specific orientations. In experiments, it was also reported that the weak tuning of compound thalamic inputs is amplified by cortical circuits (***Lien and Scanziani, 2013***), resulting in orientation-specific responses of L4 neurons. Orientation tuning of compound thalamic inputs was reported for the amplitude of temporal oscillations (F1) but not the mean firing rate (F0). It has been suggested that the tuning of the F1 amplitude could be the consequence of a spatial offset of ON and OFF subfields. In many cases this idea predicted the preferred orientation (PO) of a neuron quite well, and thus spatial segregation of subfields was proposed as a general mechanism to induce orientation selectivity in V1 (***Lien and Scanziani, 2013***; ***Pattadkal et al., 2018***; ***Jin et al., 2011***; ***Clay Reid and Alonso, 1995***). However, this concept could not explain how the orientation tuning in the amplitude (F1) of thalamic input was transformed into the output tuning in the average firing rate (F0) of V1 neurons. Moreover, the segregation of ON and OFF subfields alone could not account for the observed sensitivity of OS to spatial frequency of the grating.

The new model presented here addresses both aspects simultaneously and thus provides an integrated explanation for several hitherto unexplained features of emergent orientation selectivity in V1. The idea is that OS arises from random projections at the thalamocortical interface, exploiting the nonlinear transfer of V1 neurons. Provided the number of projections is small, a weak bias of thalamic inputs emerges by random symmetry breaking, strong enough to be amplified by the cortical circuit with the help of recurrent inhibition. We demonstrate the feasibility of such a scenario by adopting the inhibition-dominated random network described by ***Brunel (2000)*** as a model for V1, similar to previous work (***Hansel and van Vreeswijk, 2012***; ***Sadeh et al., 2014***). The neurons in this V1 network are driven by convergent inputs from untuned excitatory dLGN neurons, balanced by feedforward inhibition, and exhibit pronounced contrast-invariant tuning. Consistent with experimental observations (***Lien and Scanziani, 2013***), in our model the amplitude of the compound thalamic input (F1) converging to a V1 neuron has a weak but significant orientation bias, while the mean (F0) is insensitive to stimulus orientation. We then show that the orientation bias in F1 amplitude of the input can be transformed into a tuning for the mean firing rate (F0) of the response, exploiting the non-linear properties of single neurons. Previous computational models also studied the emergence of OS from random connectivity and the dependence on the spatial properties of the stimulus, as described in experiments (***Von der Malsburg, 1973***; ***Pattadkal et al., 2018***). However, these models were so far not able to outline any key neuronal mechanism for these phenomena.

Using numerical simulations supported by analytical considerations, we then investigate the underlying mechanism of the thalamocortical transfer. We use conventional methods of extracting ON and OFF subfields and found receptive fields that are comparable to experimental works. We also found that the input-output transformation of the orientation bias (F1 to F0) requires a nonlinear transformation. Furthermore, the contrast-invariant tuning curves of V1 neurons depend on the number of convergent thalamic inputs, as well as the spatial frequency of the grating used for stimulation. Remarkably, the model exhibits not only biologically plausible behaviour of the neuronal network, but it also explains how orientation tuning in the input is transformed into the output. This nonlinear input-output transformation is also applicable to computations in other sensory systems that rely on the processing of oscillatory signals.

## Results

To identify the underlying neuronal circuit mechanisms of orientation selective neuronal responses in V1, we performed numerical simulations of a thalamocortical network model, using sinusoidal drifting gratings for visual stimulation (cf. ***Equation 5***). The stimuli were presented at 12 different orientations, uniformly sampling all orientations between 0° and 180° in discrete steps of 15°. The movement direction was always orthogonal to the orientation of the grating.

In order to directly compare our results with experiments and other models, we also stimulated the network with flashed sparse noise arrays to estimate the receptive fields of neurons.

### Orientation tuning of compound thalamic activity

In our model, neurons in the dLGN were assumed to have circular receptive fields. The activity induced in retinal ganglion cells by a grating passing by is an oscillation with the temporal frequency of the grating (cf. ***Equation 6***). Information about the orientation of the stimulus lies only in the phase of the oscillatory activation. Neither the temporal mean, nor the oscillation amplitude of single neuron activity is sensitive for the orientation of the stimulus. The actual input to cortical neurons, however, comes from multiple thalamic neurons. Here we assume that each cortical neuron receives the same number 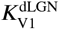 of thalamic inputs. The compound signal is again a harmonic oscillation of the same frequency, but the phases of its components matter. Depending on the relative positions of all contributing dLGN neurons, the oscillation amplitude of the compound signal may now be tuned to the orientation of the grating.

On the level of the membrane potential, the compound thalamic input to a single V1 neuron *i* is a linear sum of the responses of all presynaptic neurons

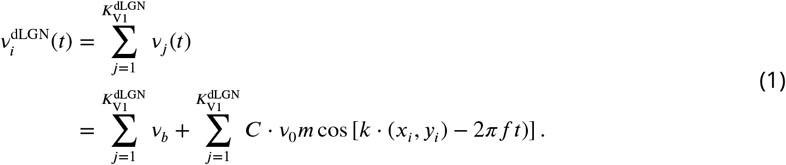

Therefore, the compound signal is again a harmonic oscillation. Its mean (F0 component) and its amplitude (F1 component) can be calculated using the Fourier theorem (***Waldert et al., 2009***)

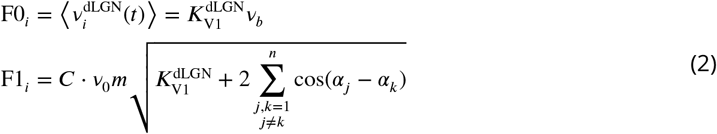

where *α*_*j*_ = *k* ⋅ (*x*_*j*_, *y*_*j*_) with *k* determined by the stimulus orientation *θ* and spatial frequency *λ* (see Methods and Materials).

In our model, the receptive field centers of dLGN neurons are randomly positioned, assuming a uniform coverage of the visual field. Each V1 neuron receives input from those dLGN neurons whose receptive fields are closest to its receptive field center. As a result, the receptive fields of V1 neurons are not all of the same size, although the number of inputs is the same (see ***Figure 1***). ON center and OFF center dLGN cells are randomly mixed. Importantly, the mean (F0 component) of the compound thalamic input does not depend on the orientation of the stimulus, while the amplitude (F1 component) is significantly tuned to orientation (***Figure 1B***). It was also shown in experiments (***Lien and Scanziani, 2013***) that it is mainly the F1 component of thalamic excitation that is tuned to stimulus orientation.

**Figure 1.**
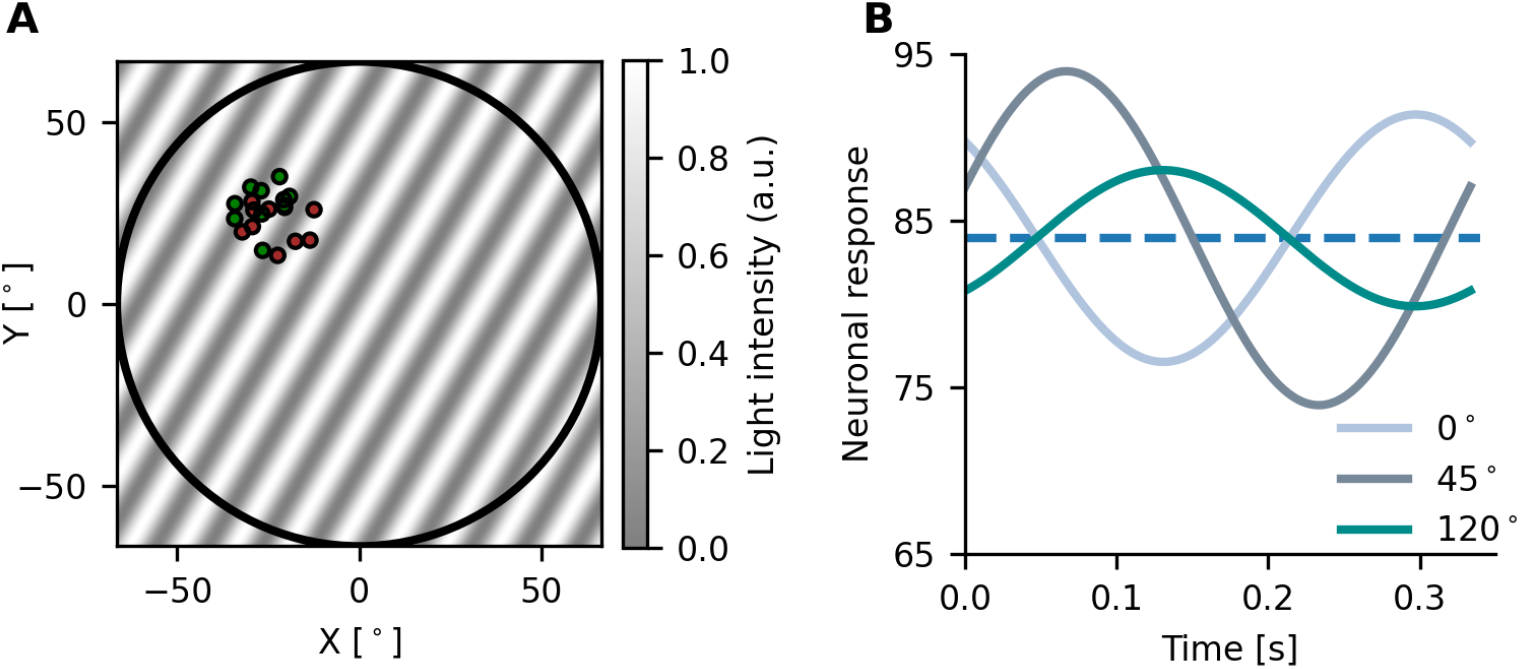
Compound thalamic input to the cortex. **A** In the example shown here, each cortical neuron receives input from 100 dLGN neurons. The receptive field of each dLGN neuron is indicated by a small circle, shown are 20 out of 100 receptive fields. Red and blue color denotes ON and OFF center receptive fields, respectively. Receptive field centers are randomly distributed and uniformly cover the (larger) receptive field of the target V1 neuron. The background shows a stimulus grating with an orientation of 60° and a spatial frequency of 0.08 cycles per degree (cpd). **B** Not only the activity of individual dLGN neurons, but also the compound signal of a group of dLGN neurons reflect the temporal modulation induced by the drifting grating stimulus. Solid lines of different colors correspond to the temporal responses for different orientations of the grating, respectively. The dashed line indicates the temporal mean of the compound signal, which does not depend on stimulus orientation.

### Nonlinear signal transmission of single LIF neurons

We have demonstrated that in our model the oscillation amplitude of thalamic compound input to cortical neurons is tuned to stimulus orientation. As the temporal mean of the compound input is untuned, it is necessary to explain how information about orientation in the input is transformed to a tuned spike response of the neuron. To derive a quantitative description of this transformation, we assume that a LIF neuron receives effective excitatory and inhibitory input matching the input level in our network simulation. The compound excitatory input is again a harmonic oscillation, and the inhibitory input does not vary in time. Therefore, the effective input to the LIF neuron is characterized by a baseline and by the amplitude of the oscillation, the phase of which is irrelevant for the questions discussed here. As the input is realized as a Poissonian barrage of action potentials with time-varying rate, we have an effective description of the resulting postsynaptic current as Gaussian White Noise with a mean *μ*_*t*_ and a fluctuation amplitude *σ*_*t*_ that depends on the input rates (***Brunel, 2000***). For simplicity we assume that the neuron is always at its steady state, producing an output that follows the relatively slow temporal modulation of its input. Its instantaneous firing rate *v*_*t*_ is then given by the nonlinear transfer function

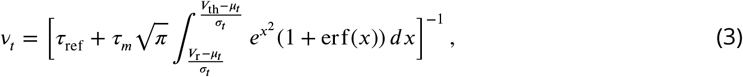

where the parameters *τ*_*m*_, *V*_th_ and *V*_r_ represent the biophysical parameters of the neuron. The mean response rate of the neuron is conceived as the temporal mean of *v*_*t*_. As the instantaneous firing rate *v*_*t*_ has the same period as the input oscillation, it is sufficient to average over one oscillation period to obtain the temporal mean (see ***Figure 2***). It is obvious that the mean of the output (***Figure 2A3***) does not correspond to the mean of the input (***Figure 2A1***) on the nonlinear transfer curve (***Figure 2A2***). In other words, the mean input is not the only factor that contributes to the mean output. When the operating point is in the nonlinear regime, the oscillatory input curve is distorted by the transfer function and the amplitude of the input oscillation also contributes to the mean response. The stationary rate model (SRM) is normally used to describe the input-output relation of a LIF neuron for stationary Poisson input. As in the present scenario the input is not stationary, however, we have to additionally account for the lowpass filter properties of the postsynaptic membrane. This motivates the dynamic rate model (DRM) adopted here. Since the excitatory input is oscillatory, its amplitude is attenuated by the frequency-dependent factor 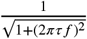 that can be derived by Fourier transforming the leaky integrator equation for the subthreshold response of the neuronal membrane. We compared the output firing rates of both firing rate models (SRM and DRM) and the simulated LIF model (SIM) for different input frequencies. We generally found a good agreement between SRM and SIM at low frequencies and a significant discrepancy at higher frequencies. The DRM, in contrast, fits quite well to the SIM for all frequencies (***Figure 2***).

**Figure 2.**
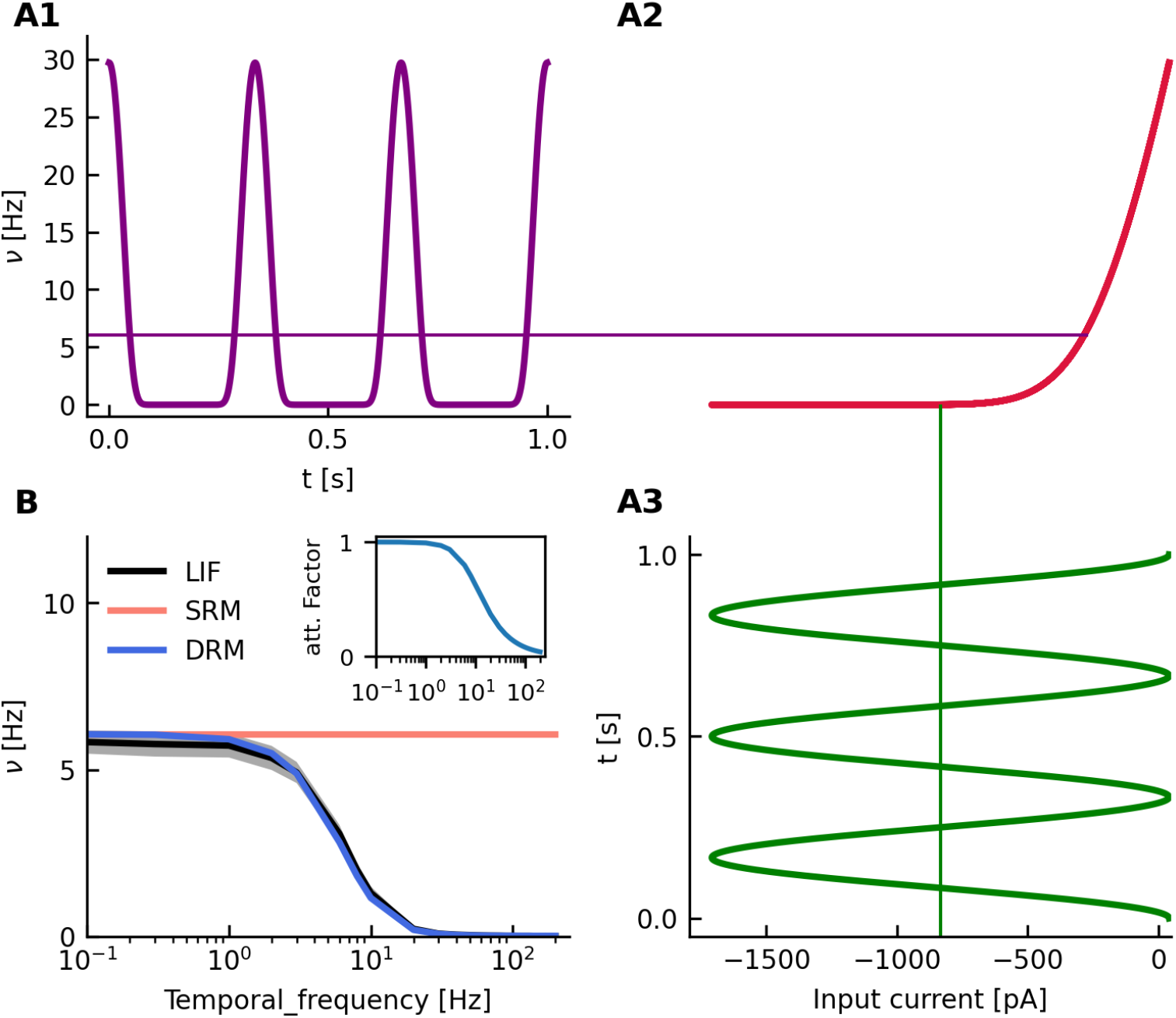
Nonlinear signal transmission of single neurons. **A1** Time-dependent output firing rate of a single spiking neuron (leaky integrate-and-fire, LIF) during stimulation with a drifting grating. Thin lines indicate the temporal mean of the time-dependent signal of the same color. The temporal frequency of the grating is 3 Hz throughout. **A2** Stationary input-output transfer function of the sample neuron shown. It is derived from the standard diffusion approximation. The output firing rate is scattered against the input current. **A3** Time-dependent current input to the LIF neuron, induced by a drifting grating. As it is a superposition of harmonic oscillations with a common frequency, it is again a harmonic oscillation of the same frequency. **B** Comparison of single neuron firing rates of three different models, for a wide range of temporal frequencies: numerical simulation of a spiking neuron (LIF), static rate model (SRM), dynamic rate model (DRM). The black line and gray shadow indicates the mean±std of numerical simulations of duration 60 s across 100 LIF neurons. The inset shows the change of the attenuation factor with temporal frequency.

The simulation of a LIF neuron revealed a specific dependence of the output rate on both the baseline and the amplitude of the input oscillation. Therefore, we separately investigated the effects of changing baseline and amplitude. When the baseline of the input is fixed and the oscillation amplitude increases (***Figure 3A***), the output firing rate also gets larger (***Figure 3B***). Observe that the mean output firing rate varies nonlinearly with the oscillation amplitude (***Figure 3C***). A similar non-linear dependency was obtained when fixing the oscillation amplitude and changing the baseline of the input (***Figure 3D-F***). Altogether, this implies that the output of a single neuron depends on the baseline and the amplitude of the input in a nonlinear fashion.

**Figure 3.**
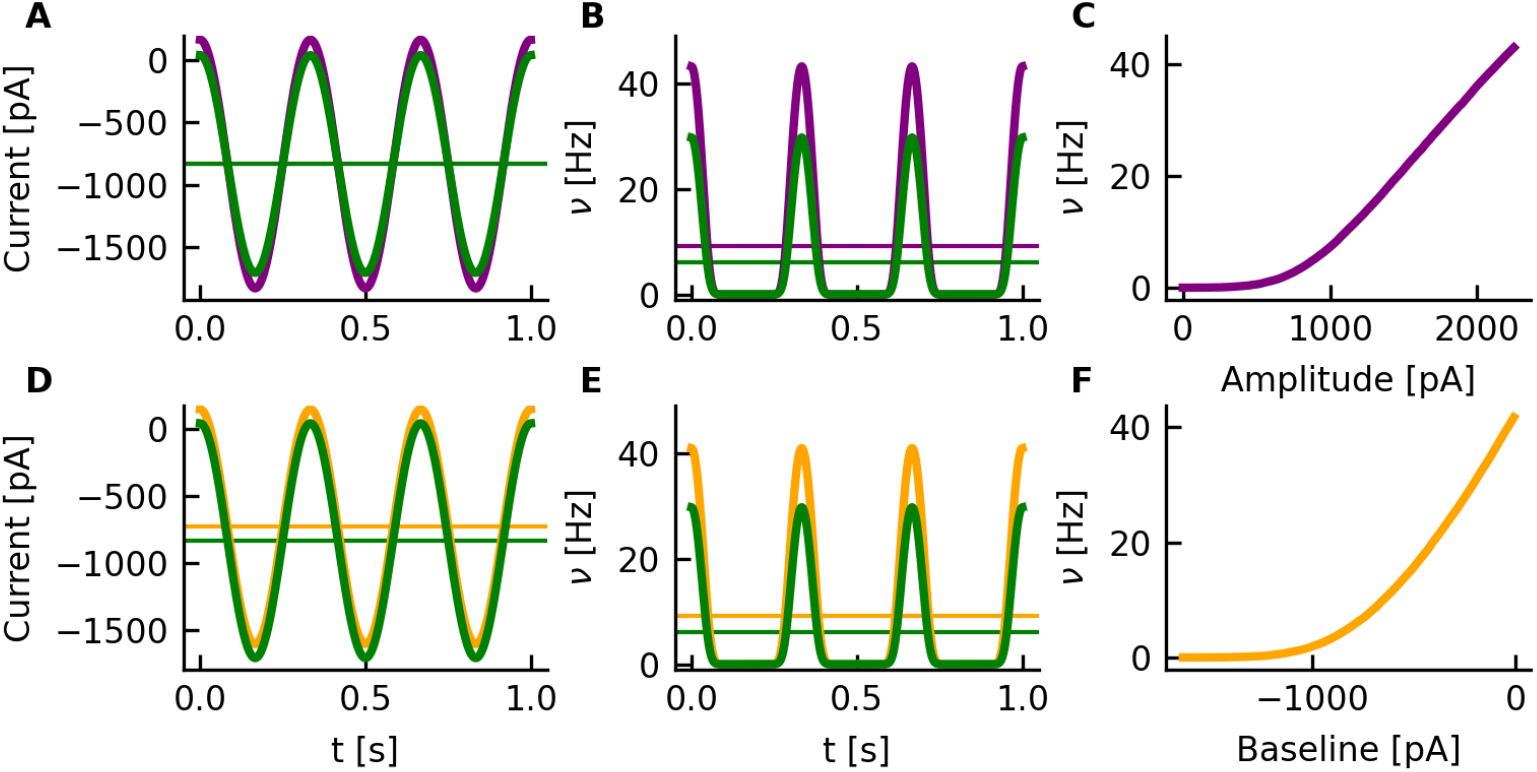
Nonlinear transduction of oscillatory input to a neuron. **A** The input current to a LIF neuron is a harmonic oscillation with a certain amplitude and additive offset. The green signal is the same as in Fig 2A3. The purple signal represents an oscillation with a larger amplitude but the same baseline. Thin horizontal lines indicate the temporal mean of the signal with matching colors. **B** Time-dependent output rate of the LIF neuron (see text for parameters), obtained by numerical simulation. Note the nonlinear distortion of the harmonic oscillation offered as input. **C** The amplitude of the input current is nonlinearly transformed to the mean output rate, assuming a fixed baseline. **D-F** Orange color indicates the outcome of a changing baseline and fixed amplitude.

### Orientation selectivity emerges from random TC projections

In previous sections, we demonstrated that the compound signal of a random sample of thalamic neurons has an orientation bias. We also showed a nonlinear dependence of single neuron responses on the F0 and F1 components of their input. Combining these two findings, we now address the question how tuning in the oscillation amplitude of compound thalamic input to cortical neurons could be transformed to tuned firing rates. To this end, we devised a thalamo-cortical network model (***Figure 4***) and performed computer simulations of its activity dynamics. The network model of V1 has been described previously ***Sadeh et al. (2014)***, based on seminal work by ***Brunel (2000)***. The V1 network model consists of *N* = 12 500 leaky integrate-and-fire (LIF) neurons, of which 80% are excitatory and 20% are inhibitory. Each V1 neuron receives input from *ϵ* = 10% of all excitatory and inhibitory neurons, the connectivity is random. Inhibitory synapses are *g* = 8 times stronger than excitatory synapses, resulting in an inhibition dominated network. A new feature of the model considered here is the feedforward inhibition (FFI), which effectively provides inhibitory thalamic input on top of the direct excitatory dLGN input. Each neuron in the recurrent network receives the same constant background input, which helps adjusting the operating point and also sets the mean response rate. All spiking network simulations were performed in NEST (***Fardet et al., 2020***).

**Figure 4.**
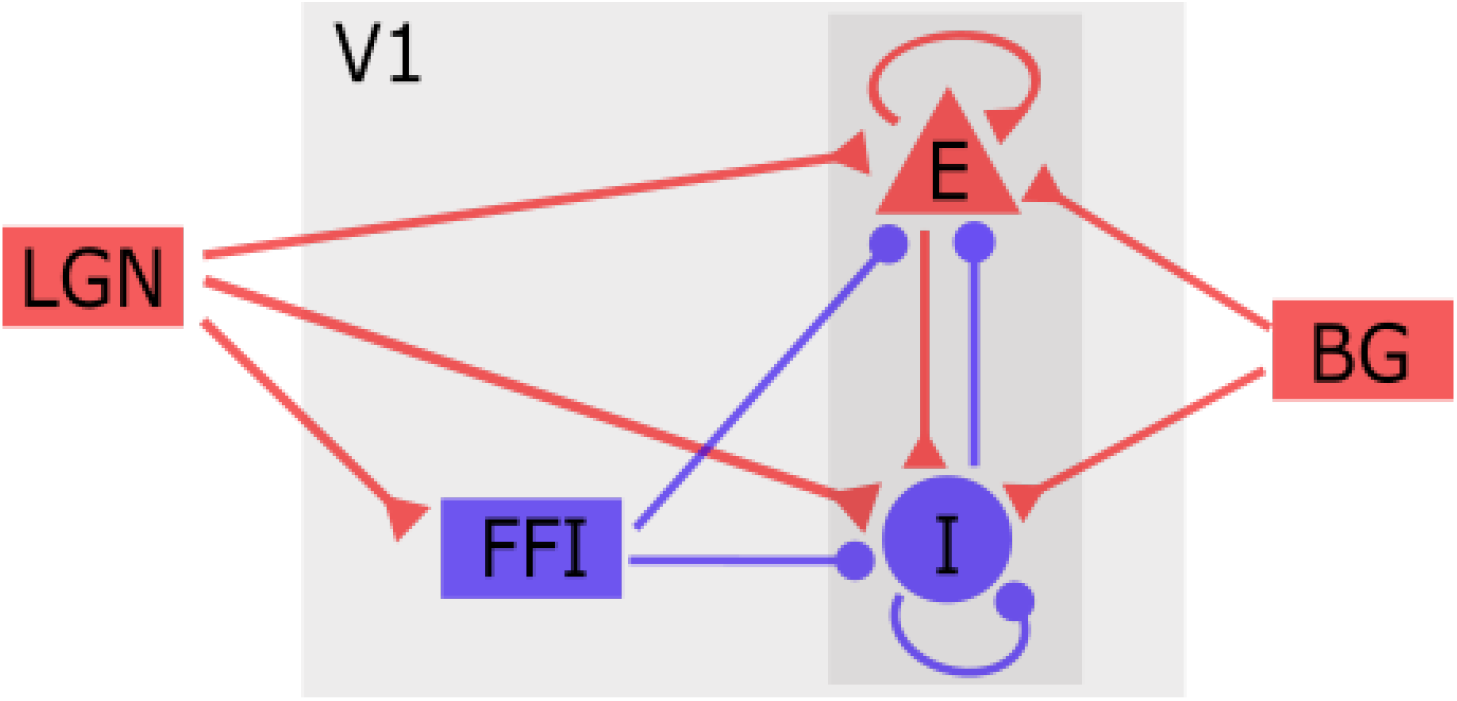
Configuration of the thalamocortical network model. The primary visual cortex (V1) comprises different types of excitatory (red) and inhibitory (blue) neurons. The thalamus (dLGN) relays light-induced activity from the retina to V1. Unspecific background input (BG) provides additional excitatory drive independent of the stimulus. Stimulus processing is collectively performed by all V1 neurons. Feedforward inhibition (FFI) and recurrent inhibition (I) together balance the recurrent excitation (E) of cortical pyramidal neurons and stabilize the operating point of the network.

In order to investigate the orientation preference of V1 neurons, we stimulated the thalamo-cortical network with sinusoidal moving gratings. We used 12 different orientations evenly distributed between 0° and 180°. Then, the tuning curves of individual recurrent V1 neurons were extracted from the recorded spike trains. To evaluate orientation preference quantitatively and study its dependence on the input, we calculated the preferred orientation (PO) and the orientation selectivity index (OSI) from the tuning curves, using methods from circular statistics. The PO is an angle between 0° and 180°. The OSI is a number between 0 and 1, where higher values denote a more pronounced orientation selectivity. First, we determined which component of the compound thalamic input conveys the orientation bias that is amplified by the V1 network. The thalamic input of a randomly selected cortical neuron is depicted in ***Figure 5***, along with some important analysis results. The input current is noisy, as a result of the random arrival of spikes generated by thalamic neurons. Fourier transformation reveals that the F0 and F1 components together carry most of the signal power. The orientation bias, however, selectively shows up in the F1 component of the compound thalamic input.

**Figure 5.**
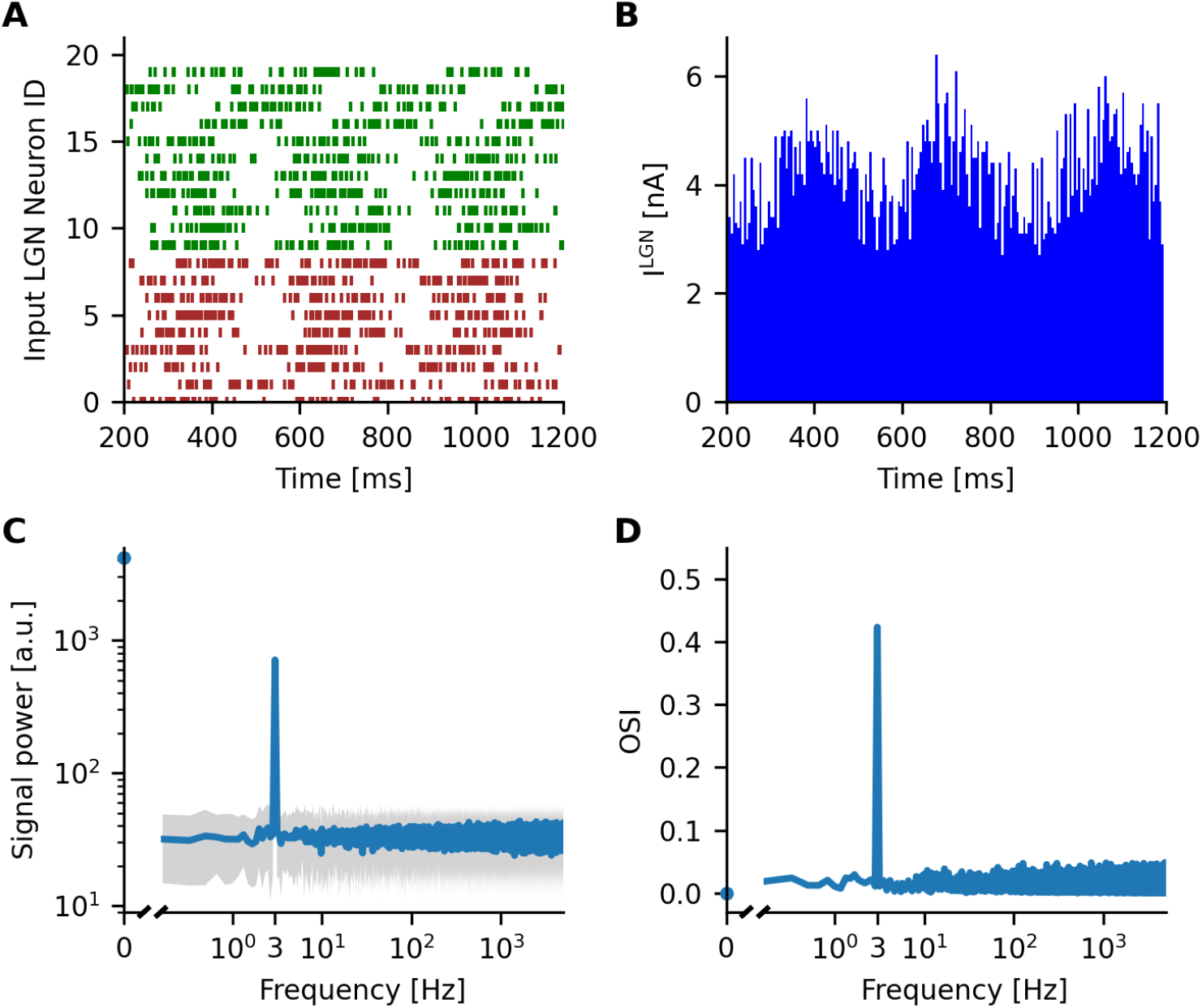
Compound thalamic inputs to a single V1 neuron. **A** Shown are the spike trains of 20 out of 100 afferent dLGN neurons that converge to a specific V1 neuron. Their locations are depicted in Fig 1A. The orientation of the drifting grating stimulus is 60°, its temporal frequency amounts to 3 Hz. **B** Compound thalamic input of all 100 afferents to a single neuron. The current is calculated from the number of spikes arriving in each time bin of the simulation (bin width 5 ms). **C** Power spectral density of the compound input signal computed by Fast Fourier Transform (FFT). The solid blue line represents the mean signal power over 50 trials, each of duration 6 s. The grey shaded area indicates the mean ± standard deviation across trials. **D** Orientation selectivity index (OSI) of the mean signal power, extracted separately for each frequency channel. Significant orientation tuning emerges only for the temporal frequency of the grating at 3 Hz.

The total feedforward input to recurrent cortical neurons is composed of time-dependent excitation from dLGN neurons, inhibition from cortical FFI neurons, and constant excitatory background input. The latter is not considered here, as it is identical for all recurrent neurons and does not convey any information about the stimulus. For each recurrent neuron, three tuning curves are extracted, namely for the mean 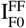 and the amplitude 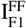 of the feedforward input current, and for the mean output firing rate *v*^V1^. All three types of tuning curves are plotted in ***Figure 6***, for a random sample of recurrent neurons. These curves essentially confirm the outcome of our single-neuron analysis. The mean input current 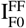 is essentially untuned, while the oscillation amplitude of 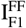 significantly varies with the orientation of the stimulus (***Figure 6***, top). The orientation bias in the input is then transformed into responses of recurrent neurons that exhibit a pronounced orientation selectivity (***Figure 6***, bottom).

**Figure 6.**
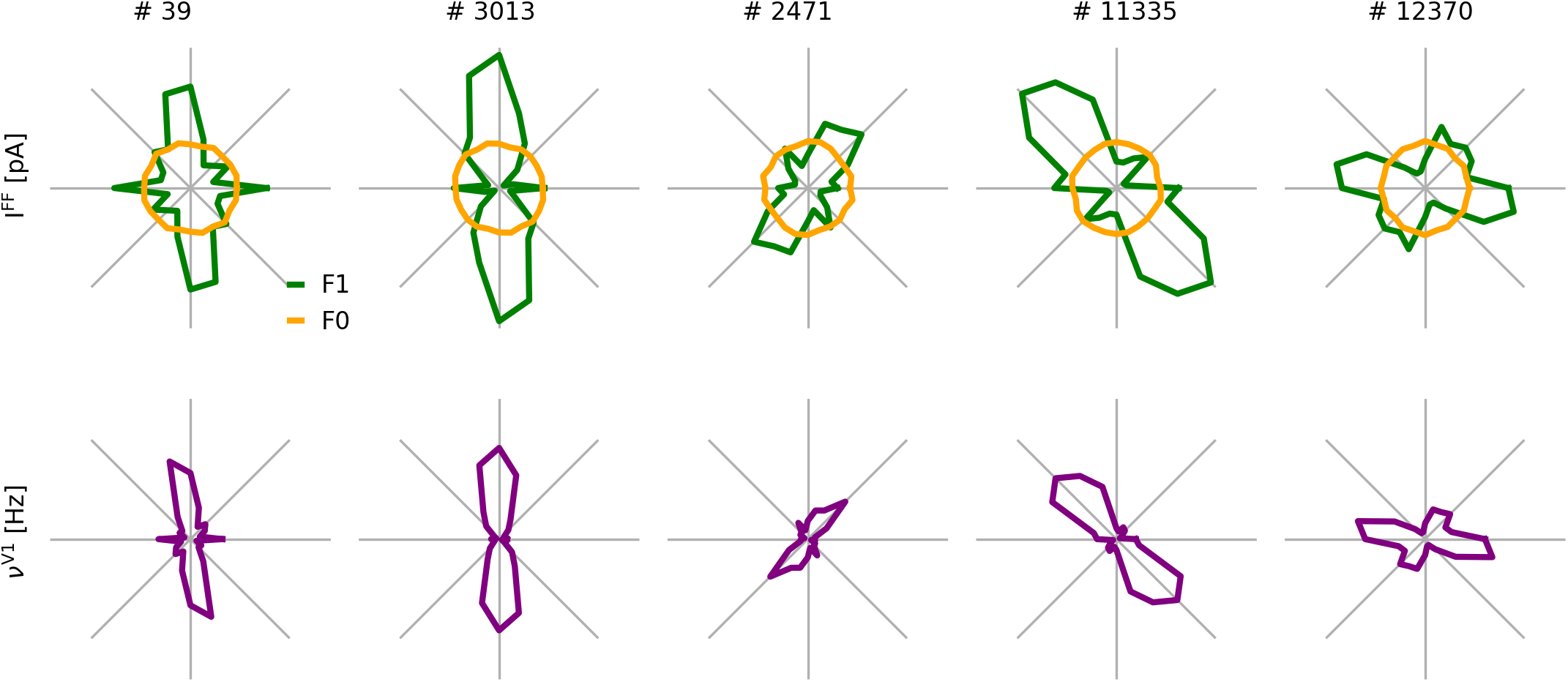
Input and output tuning curves of V1 neurons. Examples of matching input (top) and output (bottom) orientation tuning curves in polar coordinates (360°). The radial axis indicates the F0 (orange) and the F1 (green) component of the input current, as well as the mean output firing rate (purple), for five different neurons, respectively.

Next, we jointly quantified the orientation preference of cortical neurons and their correspond-ing thalamic inputs. The coefficients 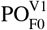 and 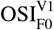 account for the PO and the OSI of the firing rate responses of recurrent V1 neurons, respectively. Since the orientation bias of the compound thalamic input shows up in F1, but not in the F0 component, its tuning is characterized by 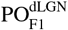 and 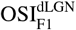, respectively. Our simulations demonstrated very clearly that orientation selectivity of V1 neurons can indeed emerge from random thalamo-cortical projections (***Figure 7B***). The OS of neuronal responses is strongly correlated with the OS of their thalamic inputs in the F1 component, which is fully consistent with experimental observations (***Li et al., 2013***; ***Lien and Scanziani, 2013***). A small residual discrepancy between input and output is due to lateral inputs from the recurrent network, as we will demonstrate later.

**Figure 7.**
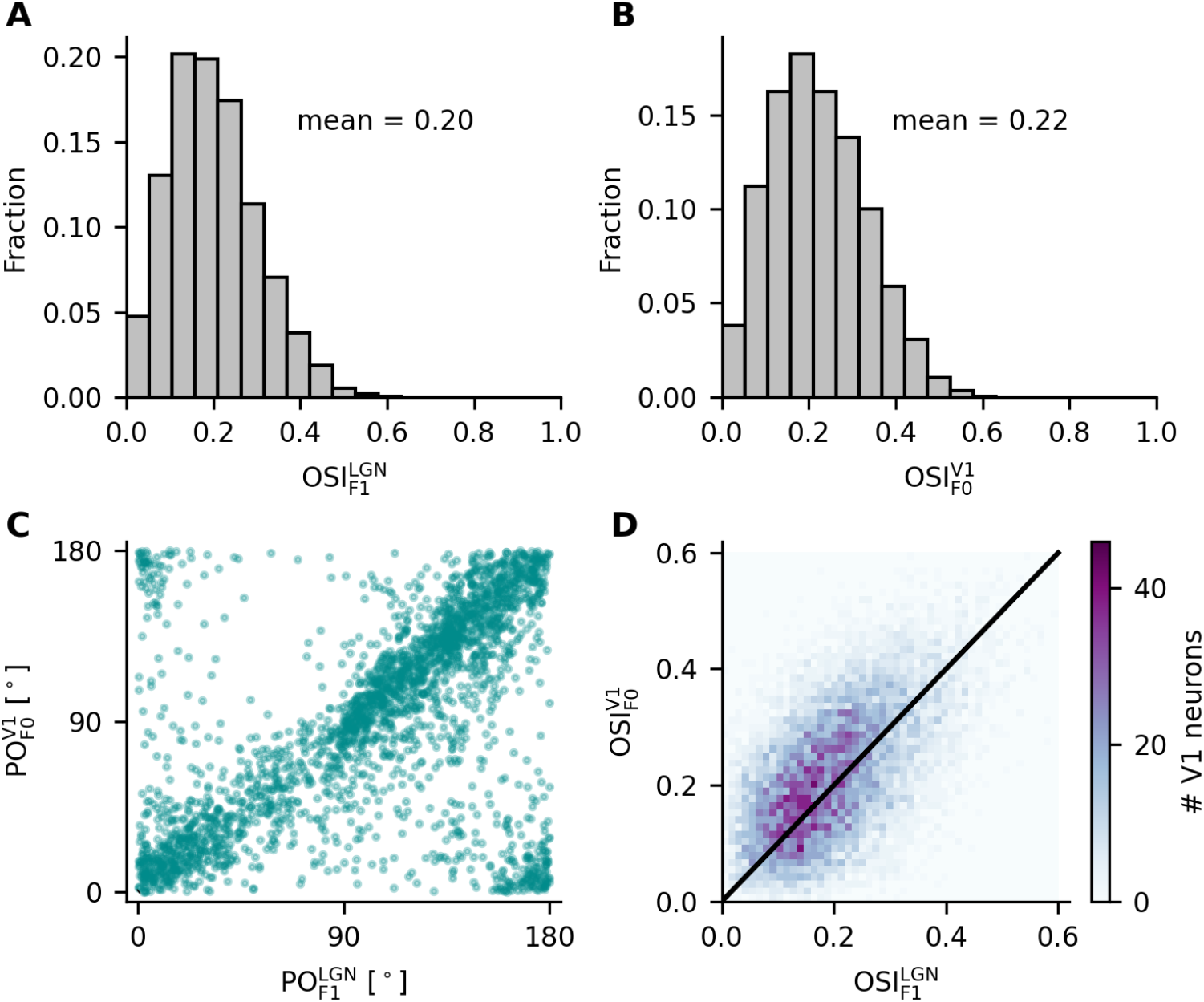
Orientation selectivity of the V1 population. Distribution of the OSI for all recurrent excitatory and inhibitory neurons in V1 (**B**) and their respective thalamic compound inputs (**A**). Unlike the output OSI, which is extracted from the orientation tuning of mean firing rates (F0), the input OSI is calculated from the oscillation amplitudes (F1) as the mean input is untuned. The scatter plots for input vs. output PO (**C**) and OSI (**D**) of all neurons show that, on average, the output OSI is slightly larger than the input OSI.

### Receptive fields in thalamus and cortex

After having shown that orientation selective responses can emerge from randomly sampling the visual field, we next investigated the receptive fields (RF) of neurons at all stages of the visual pathway represented in our model and compared them to experimental observations. As described in ***Lien and Scanziani (2013)***, we also used flashed black or white squares against a gray background as stimuli and estimated receptive fields using reverse correlation. In some neurons, the estimated PO of the RF (RF_Pref_) obtained by connecting the peaks of ON and OFF subfields was similar to the PO extracted from moving grating stimuli (Grating_Pref_) (see ***Figure 8*** #39, #3013, #11335). In other examples (#2471, #7400), however, where RF_Pref_ deviated from Grating_Pref_, the detailed shape of all subfields must be considered to predict the tuning curve. This phenomenon was also illustrated in ***Pattadkal et al. (2018)***. Since OS in V1 is essentially determined by its thalamic input, it comes as no surprise that thalamic inputs and neuronal responses have generally very similar RFs (mean correlation coefficient ≈ 0.68).

**Figure 8.**
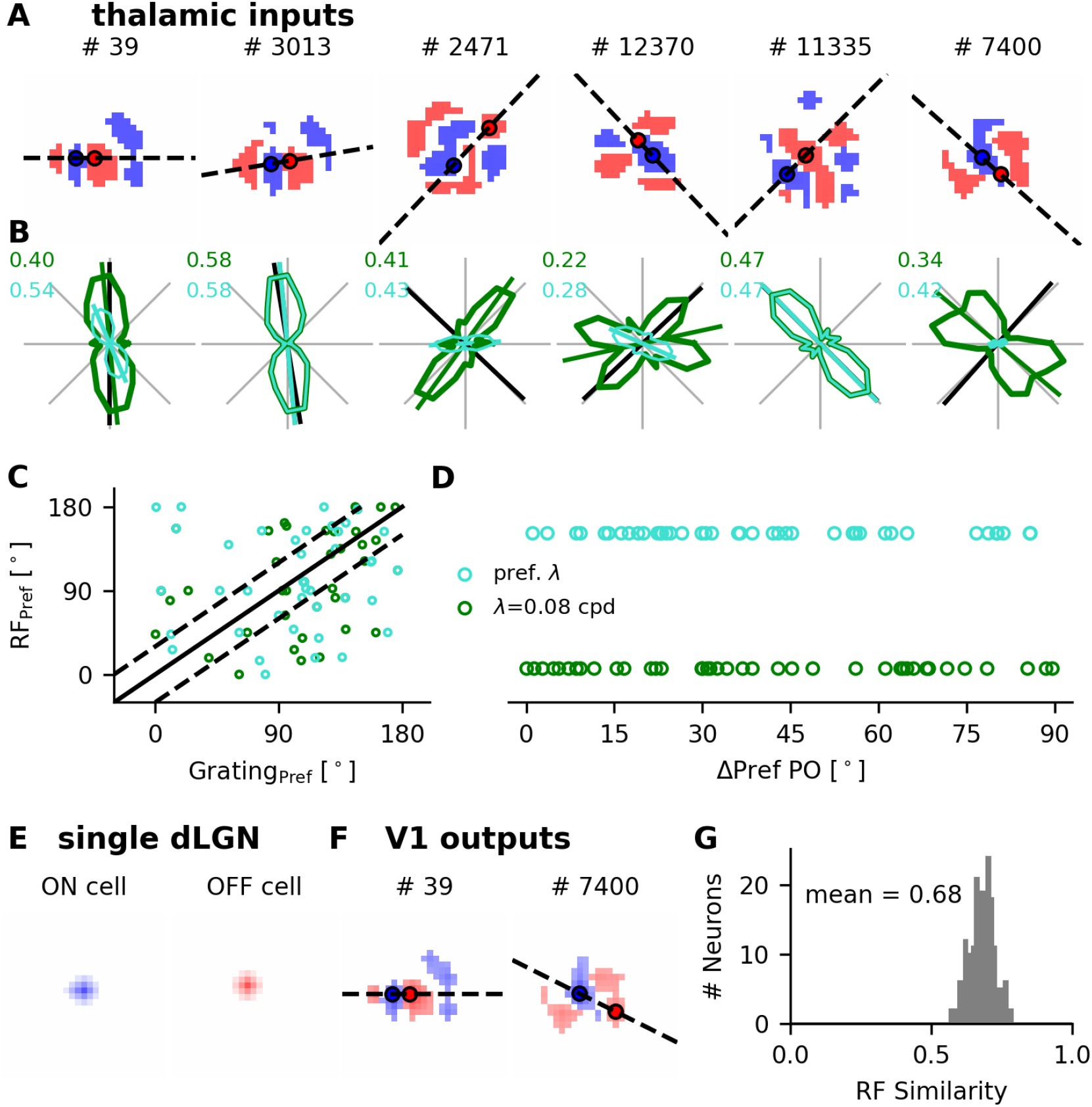
Receptive fields of recurrent V1 neurons and their thalamic inputs are similar. **A**. Receptive fields of the thalamic inputs to V1 neurons. Blue and red colors show ON and OFF subfields, respectively. The circles indicate the peaks of ON and OFF subfields, and the estimated PO is orthogonal to the line connecting the circles (dashed). **B**. Tuning curves of the thalamic inputs to the V1 neurons shown in **A**. Green curves represent the tuning curves at a fixed spatial frequency of 0.08 cpd, while the turquoise curves are the tuning curves at their respective preferred spatial frequencies. The green and turquoise lines indicate the POs extracted from the respective tuning curves (Grating_Pref_). The black lines indicate the POs estimated from their receptive fields (RF_Pref_). **C**. Scatter of Grating_Pref_ vs. RF_Pref_ for 40 neurons. The solid line is the main diagonal and the dashed lines indicate a shift by ±30°. **D**. The circular difference between RF_Pref_ and Grating_Pref_ at 0.08 cpd and the preferred spatial frequency, respectively. **E**. Receptive fields of two individual dLGN cells, one ON and one OFF center cell. **F**. Receptive fields of two sample V1 neurons. The RFs of their thalamic inputs are depicted in panel **A. G**. Histogram of RF similarities of V1 neurons and their thalamic inputs (*n* = 200, mean ≈ 0.68, median ≈ 0.68). The samples shown were obtained by spike-triggered averaging over 20 000 frames with no additional smoothing or fitting applied.

In addition, we also stimulated the random network with locally sparse noise (see Methods and Materials and ***Figure S1***). Again, the RFs of neuronal responses in V1 were very similar to the RFs of their respective thalamic input. However, the RF structure was a bit different from those obtained by flashed squares stimuli: the RFs estimated from sparse noise were generally more intricate. This was probably due to a different spatial resolution of sparse noise (0.2°) as compared to flashed squares (5°).

### Nonlinear transfer of the network

As demonstrated by numerical simulations of spiking networks, orientation selectivity in V1 can emerge from random samples of the visual field at the interface between thalamus and cortex.

Cortical neurons amplify the weak orientation bias conveyed by their compound thalamic input. A possible mechanism based on nonlinear signal transfer was explained above, but it was left to be verified that the same mechanism could also cause OS emergence in recurrent networks. To this end, we replaced spiking neurons (LIF) by an effective dynamic firing rate model (DRM). Each neuron *i* in this rate model is characterized by an explicit input-output transfer function *F*_*i*_ (***Equation 14***), see Methods and Materials for more details. As before, we used sinusoidal moving gratings with different orientations as stimuli. Each orientation was presented for a full temporal oscillation cycle at around 33 ms. The analysis was performed in 60 discrete steps per cycle. Assuming stationary responses for each step of duration 20 ms, the output firing rate was computed as a function of the respective input current. This way, we obtained the full time-locked response of input and output (***Figure 9A,C***). As before, we extracted the transfer function from these data by relating input and output in time step (***Figure 9B***). The resulting transfer curve has a characteristic form: With increasing input, the output firing rate of neurons first rises in a convex way and then enters a linear regime. The form of this curve indeed supports OS emergence, provided the operating point can be stabilized by the network.

**Figure 9.**
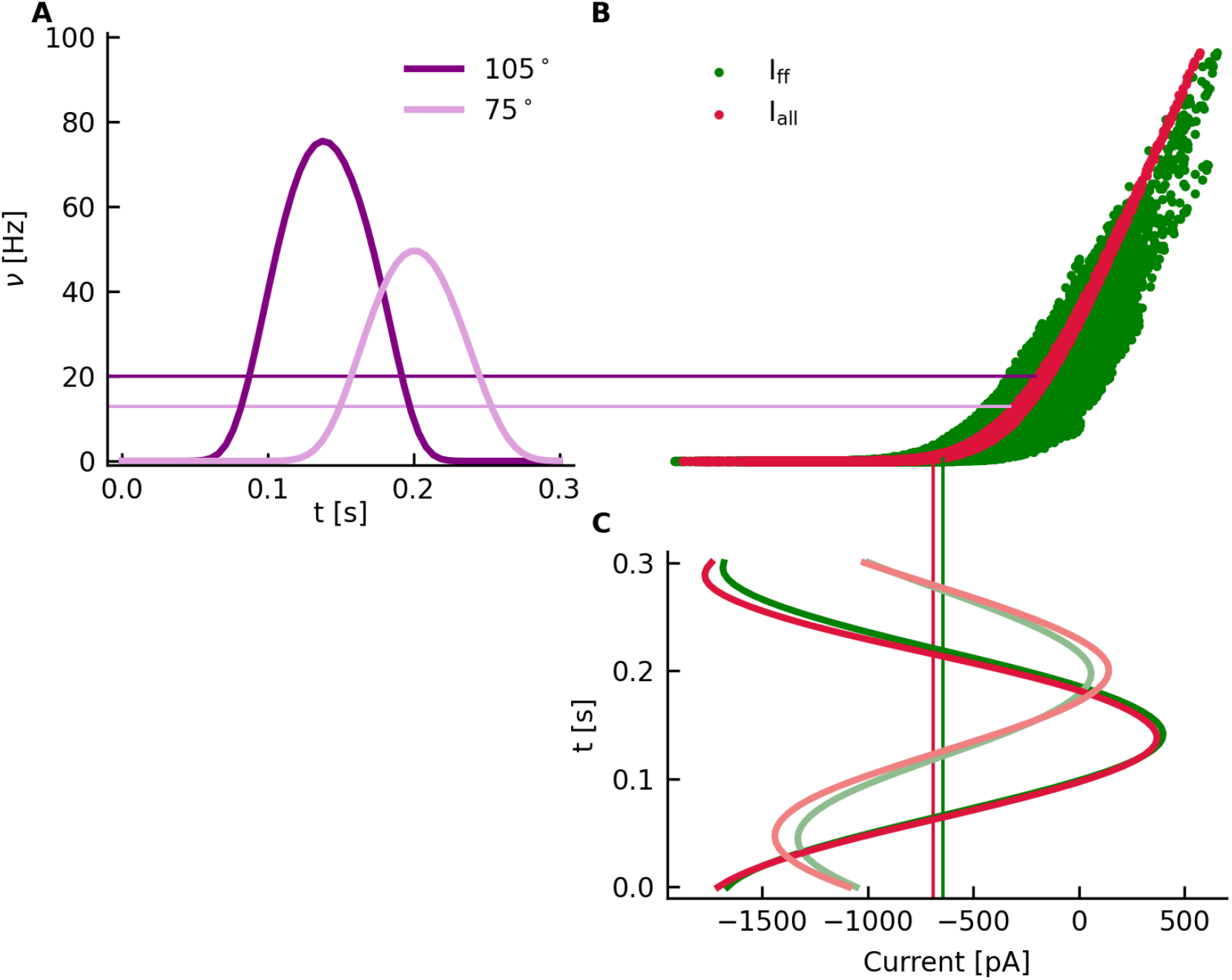
Nonlinear transfer of the network. **A** Time-dependent firing rate of a sample neuron in the recurrent V1 network. The orientation of the stimulus is 75° (lighter colors) and 105° (darker colors). Thin lines of matching colors represent the temporal mean of the signals. **B** The input-output transfer curve of a V1 neuron in the recurrent network. The output firing rate is plotted against the feedforward (green) and total including recurrent (red) input currents, respectively, as shown in **C**.

#### Effect of the recurrent input

Having identified the nonlinear input-output transfer curve of neurons in the network, we went on to characterize the individual contributions of feedforward and recurrent inputs, respectively, to the emergence of orientation selectivity. To this end, feedforward input current I_ff_ and the total input current I_all_ were calculated separately and plotted against the output firing rates of single neurons. Not surprisingly, feedforward inputs essentially follow the shape of the transfer curve, although with some uncertainty (***Figure 9B*** green dots). When the recurrent input is also taken into consideration, the relation between input and output is much more determined (***Figure 9B*** red dots). This indicates that the output firing rate is mainly caused by feedforward input and only slightly perturbed by recurrent input. This explains the strong correlation between preferred orientations of input and output, as well as the residual discrepancies between them, observed in simulations (see ***Figure 7***).

#### Comparison of spiking neurons and firing rate neurons

In the previous section, we have seen consistent behavior of simulated LIF neurons and the DRM, for a range of different temporal frequencies (***Figure 2D***). For a moving sinusoidal grating with temporal frequency *f* = 3 Hz, the DRM is able to track the dynamics of the input signal with high fidelity. Therefore, we consider it as a useful approximation of the spiking neuron. A comparison of both models at this frequency also yields good consistency with regard to the preferred orientation (PO) and orientation selectivity (OSI) of single neurons (***Figure 10***), respectively. Residual discrepancies between the two models are explained by random fluctuations in the timing of individual spikes.

**Figure 10.**
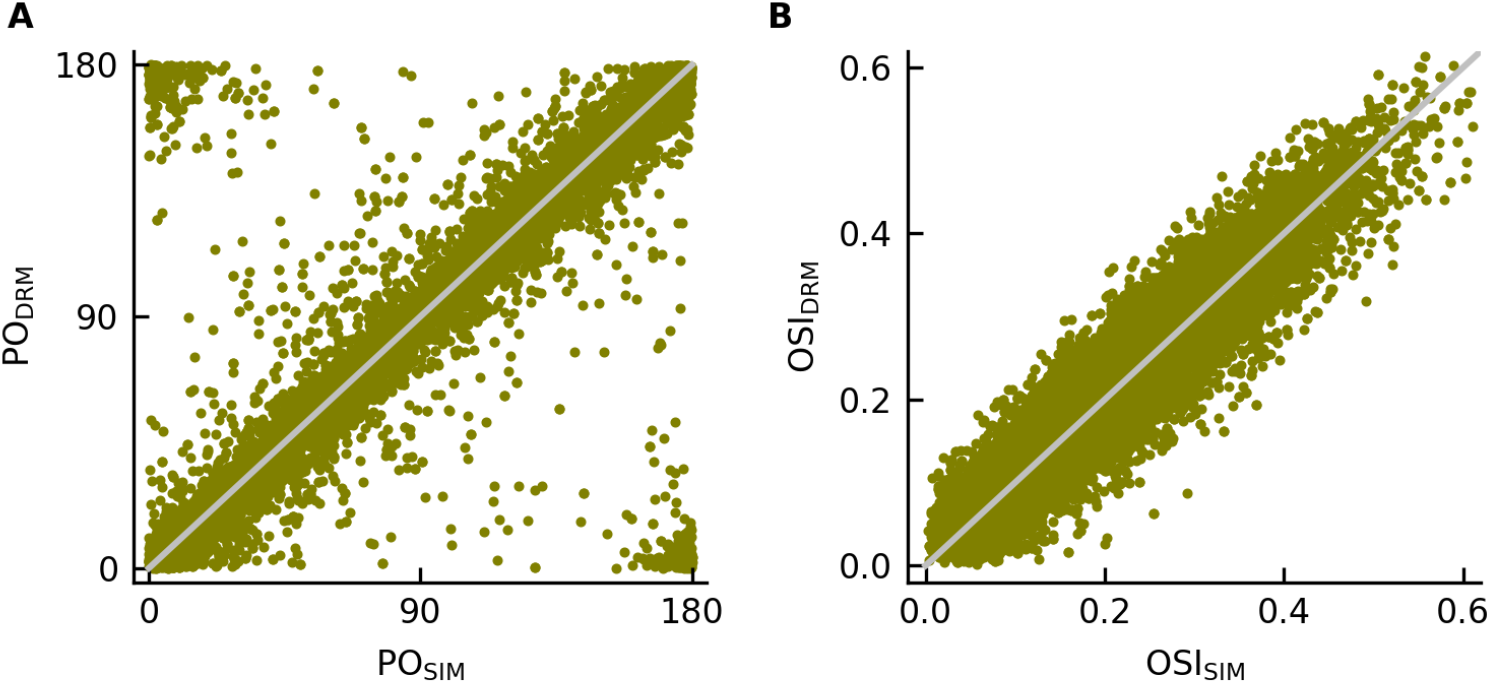
Performance of the DRM compared to numerical simulations of spiking neurons. The behavior of individual LIF neurons in network simulations is compared to predictions from the dynamic rate model (DRM), see text for details. Shown are scatter plots of the PO (**A**) and the OSI (**B**) for all recurrent V1 neurons. The gray diagonal line indicates a perfect match.

### Parameter dependence of orientation selectivity

In the previous paragraph, we have outlined a candidate mechanism how orientation selective responses of recurrent V1 neurons can emerge from random thalamocortical connectivity. Employing a nonlinear transfer function, the F1 tuning in the input is transformed into a F0 tuning curve at the output. As a result, the input F1 component is an essential determinant of the output OS in recurrent V1 neurons. As expressed by ***Equation 2***, the F1 component of the input depends explicitly on the number of dLGN neurons that converge on a recurrent V1 neuron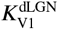. In addition, it depends on the spatial frequency of the moving grating *λ*. In this section, we investigated the detailed dependence of the output OS on these two parameters.

#### Thalamo-cortical convergence number affects neuronal selectivity

To elucidate the role of the number of thalamo-cortical projections 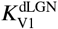 for orientation selectivity of cortical neurons, we determined its impact on the output OSI of recurrent V1 neurons. Throughout all simulations, the size of thalamic receptive fields and the mean current 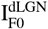 corresponding to compound thalamic input to V1 neurons were kept fixed. Mean and standard deviation of the OSI across all neurons are depicted for different values of the thalamo-cortical convergence number (***Figure 11***). We found that strong and reliable tuning is obtained for a very broad range of values for 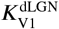, covering more than two orders of magnitude between a few and a few hundreds (***Figure 11B***). Anatomical counts, in fact, yielded numbers in the range between 15 and 125, depending on the animal species (***Alonso et al., 2001***; ***Peters and Payne, 1993***; ***Potjans and Diesmann, 2012***). If not stated otherwise, the convergence number in our model was set to 100.

**Figure 11.**
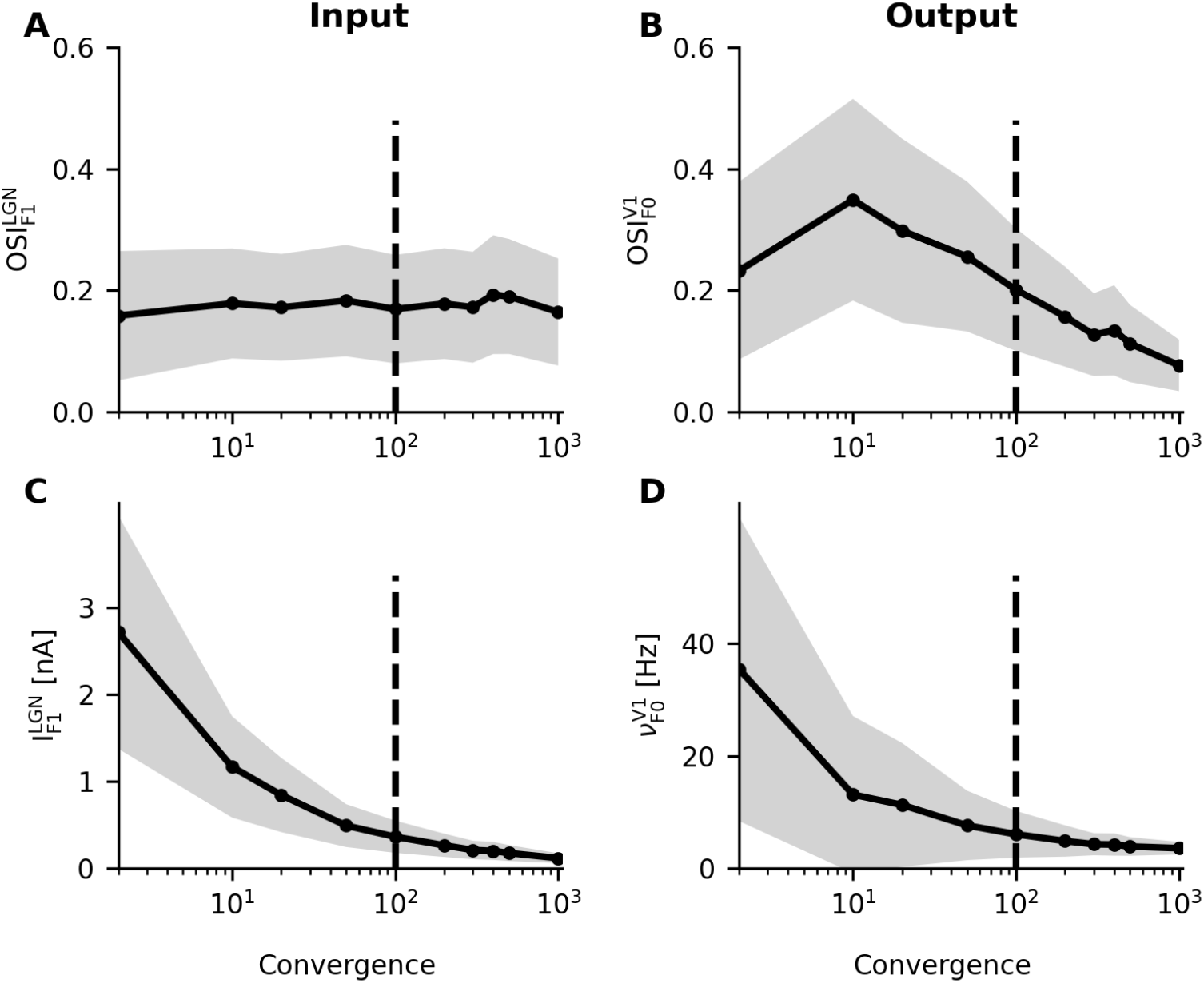
Thalamo-cortical convergence and orientation selectivity. Four quantities are plotted against the number of thalamic afferents converging on a single cortical neuron: **A** The OSI of oscillation amplitudes of the compound thalamic input, **B** the OSI of firing rate output in recurrent neurons, **C** the amplitude of compound thalamic input current oscillations, and **D** the mean firing rate of recurrent neurons. Solid lines and gray shaded areas represent the mean ± standard deviation. Depending on the animal species, convergence numbers between 80 and 200 have been reported. A convergence number of 100 was chosen in most of our simulations (dashed line).

We also investigated the dependence of the F1 component of compound thalamic input on the number of convergent inputs, as it represents the most important determinant of the output orientation preference. For increasing convergence numbers, the oscillation amplitude of the compound thalamic input current 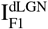 decreases (***Figure 11C***), while the OSI of the input amplitude remains at a fixed level (***Figure 11A***). As a result of nonlinear signal transfer, the output OSI depends on the convergence number in a complex manner (***Figure 11B***). When the convergence number is low, the oscillation amplitude is large, resulting in high output firing rate. In this case, the operating point is almost shifted outside the nonlinear range, and the output OSI becomes smaller. When the convergence number gets larger, the input OSI remains unchanged, but the oscillation amplitude decreases. Therefore, the output OSI declines as the oscillation is amplified less. Overall, the orientation selectivity of the output does not only depend on the oscillation amplitude of the thalamic input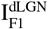, but also its orientation selectivity 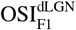.

#### Spatial frequency affects neuronal orientation tuning

The second parameter that affects the orientation preference in the compound thalamic input and further in V1 is spatial frequency (see ***Equation 2***). Sinusoidal moving gratings at different spatial frequencies ranging from 0.001 cpd to 0.4 cpd were used as visual stimuli. Orientation preference (PO and OSI) was extracted from the single-neuron tuning curves at each spatial frequency. For the network layout considered in our model, strongest tuning was observed between 0.06 cpd and 0.08 cpd, and the tuning became rather weak for very small (below 0.01 cpd) and for very large (above 0.3 cpd) spatial frequencies (***Figure 12A***). This showed that the strength of orientation tuning (OSI) of the output was strongly affected by the spatial frequency of the stimulus, and the strongest tuning was obtained for a spatial frequency at about 0.08 cpd.

**Figure 12.**
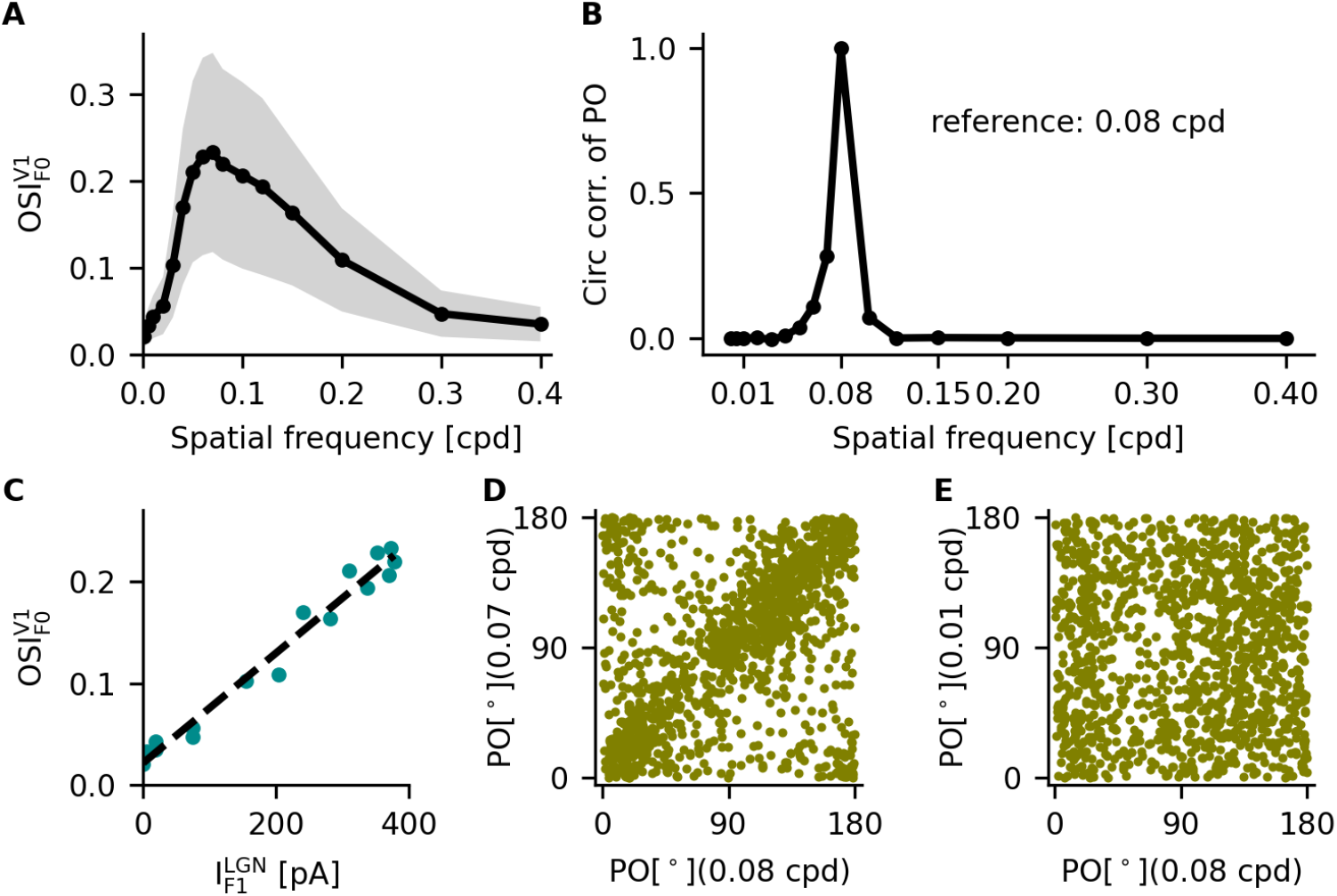
Impact of spatial frequency on orientation selectivity. **A** The OSI depends on the spatial frequency of the drifting grating used for stimulation in the model. The solid curve and the gray shaded area represent the mean ± standard deviation. **B** The PO of individual neurons in the same network changes with the spatial frequency of the grating. The PO at 0.08 cpd is very different from the PO for deviating spatial frequencies, as indicated by the circular correlation coefficient. **C** The oscillation amplitude of compound input current is the most important determinant for the OSI of the output firing rate in recurrent neurons. **D**,**E** A similar picture emerges by directly comparing the PO of all recurrent neurons at nearby spatial frequencies (0.07 vs. 0.08 cpd) and at strongly deviating spatial frequencies (0.01 vs. 0.08 cpd).

In addition, we observed that the preferred orientation (PO) of single neurons was different for different spatial frequencies. To quantify the changes in PO, we first determined the PO for different spatial frequencies, for all neurons in the network. Separately for each spatial frequency, we then calculated the circular correlation (see Methods and Materials) of these angular variables with the PO obtained for the same neuron at the reference spatial frequency of 0.08 cpd. The correlation is very high for similar frequencies and very small for distant frequencies (***Figure 12B***). The correlation coefficient is around 0.35 for 0.07 cpd (***Figure 12D***), while it is close to 0 for 0.01 cpd (***Figure 12E***). Similar observations were also made in experiments in rodents and higher mammals. The spatial frequency has generally a strong impact on the OSI, and different spatial frequencies lead to a different PO in single neurons (***Ayzenshtat et al., 2016***; ***Pattadkal et al., 2018***). Note that models of primary visual processing that link orientation selectivity with excitatory and inhibitory subfields of neuronal receptive fields cannot explain such dependencies in principle, as the spatial frequency of the stimulus is not taken into account.

### Contrast-invariant orientation tuning

The perceived orientation of a stimulus should, ideally, not depend on the stimulus contrast. Contrast-invariant tuning curves were indeed widely observed in the visual cortex of cats as well as mice (***Ferster and Miller, 2000***; ***Priebe and Ferster, 2008***; ***Niell and Stryker, 2008***). Here, we report that contrast-invariance of orientation tuning is also a property of neuronal responses in our model. Sinusoidal drifting gratings with contrasts varying between 0 and 1 (see Methods and Materials) were presented for 12 different orientations, as described before. Note that the mean luminosity of the grating remained unchanged for the different contrasts considered here. As a consequence, the mean firing rates of dLGN neurons were unchanged as well, while the stimulus contrast was reflected by the amplitude of temporal oscillations. For zero contrast, the sinusoidal “grating” is just a uniform gray with the same (mean) luminosity everywhere. Clearly, no stimulus orientation can be observed under this condition. Although the response amplitudes at the preferred orientation are higher for stronger contrasts, the shape of the tuning curves does not depend on contrast (examples see ***Figure 13C,D***).

**Figure 13.**
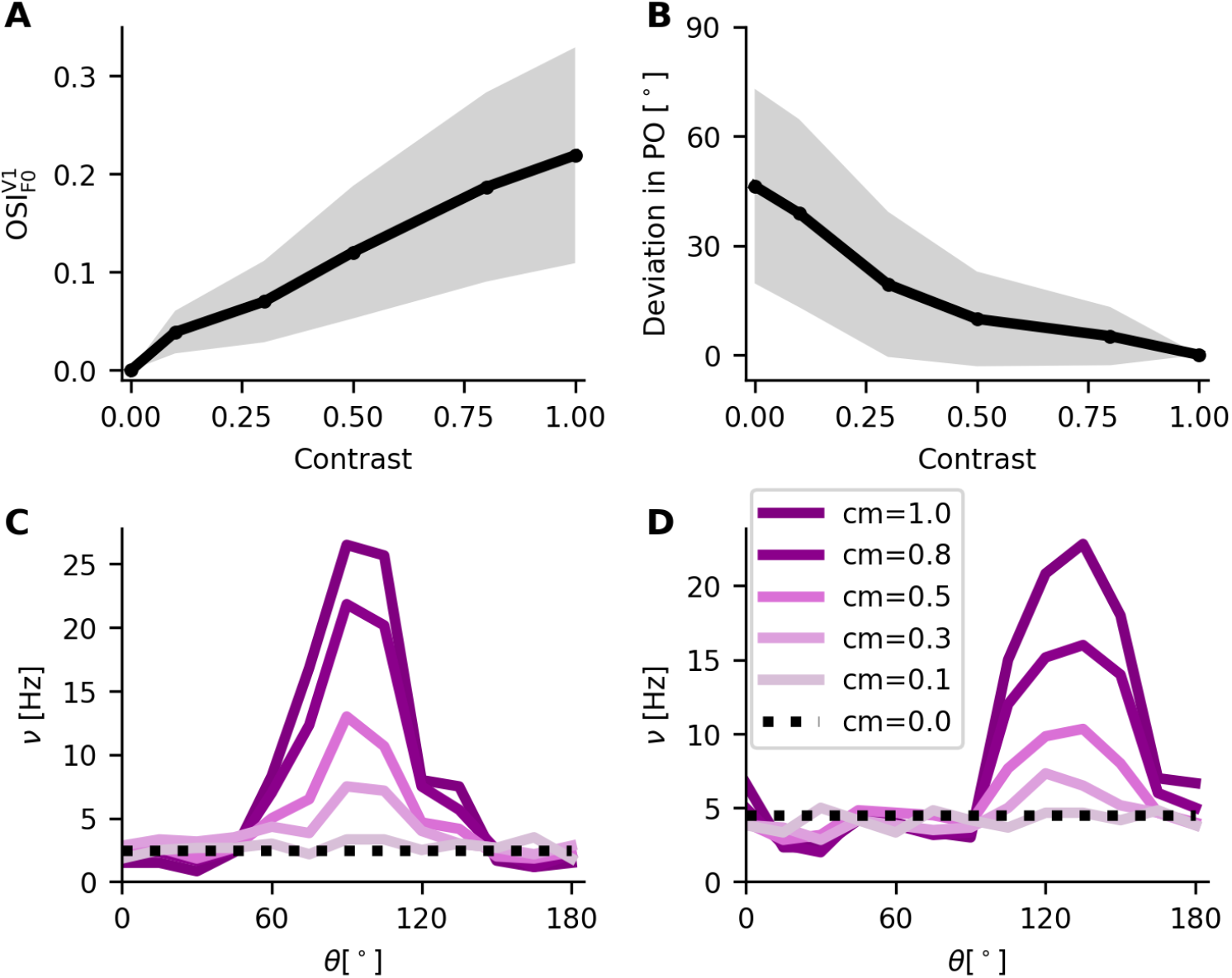
Contrast-invariance of tuning curves. **A** Shown is the OSI of V1 neurons for different values of the stimulus contrast (mean ± standard deviation). **B** Deviation of the PO in degrees for a stimulus of reduced contrast as compared to maximal contrast (1). **C**,**D** Sample tuning curves for one excitatory (left) and one inhibitory (right) neuron, at different contrasts of the stimulus. Lighter colors represent lower contrasts. Dotted lines indicate the neuronal responses at 0 contrast.

To investigate the impact of stimulus contrast on orientation preference, the PO and the OSI of single neurons were extracted. We found that the OSI was generally proportional to contrast (***Figure 13A***). This reflects the fact that the tuning strength is related to the signal-to-noise ratio, which is here determined by the amplitude of temporal oscillations and the offset. We also quantified the stability of tuning by calculating the absolute PO difference for single neurons at reduced contrasts compared to the maximum contrast 1. ***Figure 13B*** demonstrates that the PO extracted from simulated spike trains is rather stable. For very low contrasts, however, low signal-to-noise ratios do not support reliable estimates of the PO. Our analysis supports the notion that orientation tuning curves are contrast-invariant, consistent with what has been reported from experiments.

## Discussion

We studied the mechanism and the properties of emergent orientation selectivity in the early visual system. In fact, our analysis of the thalamocortical pathway combined different perspectives:

We used numerical simulations to demonstrate that orientation selective responses in the pri-mary visual cortex can emerge from random sampling the visual field based on unstructured projections from the thalamus to cortex. No matter whether the stimulus consisted of moving gratings, flashed squares or sparse noise, we found that the estimated PO was linked with the segregation and the intricate shape of the ON and OFF subfields (***Figure 8***). In all cases, the properties of cortical responses were strongly correlated with the properties of the thalamic input. We generally found that the contrast-invariant tuning curves in V1 neurons were quite sensitive to the spatial frequency of the stimulus. Both the OSI and PO of neuronal responses were strongly influenced by the spatial frequency of the grating used for stimulation, similar to what has been found in experiments (***Ferster and Miller, 2000***; ***Ayzenshtat et al., 2016***).

The number of thalamo-cortical afferents was identified as a critical anatomical parameter of the system. Numerical simulations of our model revealed that the orientation selectivity of the output depended strongly on the number of dLGN afferents, matching the numbers known from different animal species (***Alonso et al., 2001***). All these insights combined allowed us to study the feedforward transfer of feature selectivity underlying OS emergence using analytical tools. We found that nonlinear signal transduction and input statistics together can explain the F1-to-F0 input-output transformation, as well as the strong correspondence of the PO in input and output. Both facts have been reported in experiments ***Lien and Scanziani (2013)***.

It was claimed in ***Pattadkal et al. (2018)*** that the OSI is robust to the number of convergent afferents and the spatial frequency of the stimulus. Our conclusions strongly deviate from this, as we covered a wider range of parameters. For the convergence number, we considered a range between 2 and 1 000, whereas Pattadkal et al. took only the small window between 25 and 100 into consideration. For the spatial frequency of the gratings, we tested values between 0.001 and 0.4 cpd, while they considered the range between 0.01 and 0.15 cpd only.

### Orientation tuning of dLGN neurons

In our model, by design, dLGN neurons respond equally to all stimulus orientations of oriented drifting gratings. Orientation selective responses of V1 neurons emerge, for the first time, at the interface between thalamus and cortex. In contrast to single dLGN neurons, the oscillation amplitude of compound thalamic inputs has a significant orientation bias. This bias in the oscillation amplitude (F1) is transformed into a bias of mean firing rates (F0). Contrast-invariant tuning curves result with the help of recurrent inhibition in the V1 network.

Since orientation selectivity was first described in cat visual cortex ***Hubel and Wiesel (1962)***, it has long been thought that individual dLGN neurons convey only untuned inputs to the visual cortex. However, recent experimental studies in mice revealed that some dLGN relay cells are somewhat orientation selective (***Scholl et al., 2013***; ***Tang et al., 2016***). These tuned dLGN cells indeed project to layer 4, the main input layer of V1 (***Sun et al., 2016***). In our network model, the mean and dispersion of orientation selectivity across cortical neurons is a bit smaller than reported in experiments (***Ko et al., 2013***; ***Niell and Stryker, 2008***). Accounting for individual thalamic inputs with orientation preference would potentially increase the tuning of the input amplitude and, therefore, also yield slightly stronger orientation selectivity in V1 neurons. This might bring our model even closer to experimental findings.

### Contribution of the input amplitude

In the model developed here, the orientation bias in the amplitude of input oscillations is transformed into orientation tuning of output firing rate, exploiting generic nonlinear properties of spiking neurons (input rectification induced by the spike threshold). The mean input, which is the same for all stimulus orientations, sets the operating point. Unfavorable combinations of parameters, however, may compromise the nonlinear transduction and attenuate the output tuning. Our analysis revealed that orientation selectivity of the output is mainly determined by the amplitude of input oscillations (***Figure 7D***, ***Figure 11, Figure 12C***). On the one hand, small oscillation amplitudes render the nonlinear transduction mechanism ineffective. On the other hand, large input amplitudes can lead to low output selectivity, if the modulation of the input amplitude is small. The magnitude 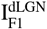 as well as the modulation 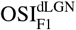 of input amplitude work together to determine the output selectivity of V1 neurons.

### Role of feedforward inhibition

To achieve optimal orientation selectivity of firing rates, it is important to stabilize the operating point in the nonlinear regime of the neuronal transfer function. In the thalamocortical network model considered here, the untuned component of the thalamic input is compensated by feedforward inhibition, and a stable F1-to-F0 transformation is enabled. Feedforward inhibition is generally associated with parvalbumin (PV) expressing GABAergic interneurons. In experiments, it has indeed been shown that PV neurons provide untuned feedforward inhibition to excitatory neurons in V1 (***Ma et al., 2010***). In addition, layer 4 PV interneurons are directly innervated by thalamocortical axons in layer 4 (***Rudy et al., 2011***; ***Ji et al., 2015***). Fast spiking basket cells, a subtype of PV interneurons, were explicitly shown to mediate the feedforward inhibition of thalamocortical inputs. These findings are consistent with the role assigned to feedforward inhibition in our network model: These interneurons are mainly driven by thalamic input, they typically fire at high rates, and their output provides essentially untuned inhibitory input to recurrent V1 neurons.

Besides feedforward inhibition in our network, there are other known pathways which might contribute to stabilize the baseline of thalamic input. For instance, the thalamic reticular nucleus (TRN) is comprised exclusivley of GABAergic interneurons and can make an indirect contribution being involved in a corticothalamic pathway. In this feedback loop, TRN cells receive input from both thalamus and cortical layer 6 and then exclusively project to thalamic nuclei. This enhanced recurrent circuit might control the excitation of thalamocortical relay cells and this way modulate thalamic signaling (***Sherman, 2011, 2016***; ***Neyer et al., 2016***; ***Coulon et al., 2009***).

### Choice of the neuron model

The emergence of orientation selectivity in our model depends in an essential way on the generic nonlinear transmission properties of spiking neurons. In our simulations, all neurons are conceived as current-based leaky integrate-and-fire (LIF) point neurons. The question arises whether cortical nerve cells in particular are well represented by this reduced neuron model. Despite the lack of structured dendrites and detailed intrinsic conductances, however, the LIF neuron model is able to capture fundamental processes performed by biological nerve cells, namely synaptic input integration and spike-based signaling. This generic model is, in fact, the most widely used model to study the dynamic behavior of large recurrent networks (***Brunel, 2000***) and has been found useful in studying information processing in neural networks (***Burkitt, 2006***). Therefore, the LIF model is a natural and adequate choice to also study the generic mechanisms underlying thalamocortical signal processing.

### Consistency with experiments

The results of our model-based analyses are widely consistent with observations reported in mouse experiments. In our model, the ON and OFF subfields of V1 neurons and their thalamic inputs were estimated from the neuronal responses to flashed square stimuli. Experimental work showed that the spatial offset of ON and OFF subfields can often predict the preferred orientation of neurons (***Jin et al., 2011***; ***Lien and Scanziani, 2013***). We found, however, that not only the offset but also the detailed shape of subfields influences orientation preference, as reported in ***Pattadkal et al. (2018)***. We can add here that the similarity between receptive fields of thalamic inputs and cortical outputs is generally quite high.

It has been proposed that the offset between the peaks of ON and OFF subfields can give rise to an orientation bias in the thalamic F1 component (***Lien and Scanziani, 2013***). A key role of visual cortex in transforming and amplifying the tuned thalamic input was also demonstrated in these experiments. In line with these findings, we also observed that the F0 component of thalamic input in our model is essentially untuned to stimulus orientation, while the F1 component has a significant orientation bias. This initial bias is then transformed into strong and contrast-invariant orientation tuning in recurrent V1 neurons. In our model, the orientation preference of V1 neuronal responses is strongly correlated with the preferred orientation of their thalamic inputs. The output has a slightly stronger orientation selectivity than the input, measured by the OSI (***Figure 7 C,D***). The orientation selectivity of recurrent V1 neurons in the model, however, is somewhat smaller than reported in experiments (***Scholl et al., 2013***; ***Pattadkal et al., 2018***). On the one hand, tuned input from the thalamus, which is not considered in our model, can potentially increase cortical orientation selectivity. On the other hand, thalamocortical projections in animals are not as random as assumed in our model. Nonrandom spatial sampling of TC projections will typically also enhance the orientation selectivity of cortical neurons. The spatial frequency of the visual stimulus has a strong impact on the preferred orientation as well as the strength of the orientation selectivity, which also has been reported in experiments (***Ayzenshtat et al., 2016***).

### Comparison with alternative models

Most previous theoretical works account of orientation selectivity referred to recordings from cats and primates (***Von der Malsburg, 1973***; ***Soodak, 1987***; ***Ringach, 2004***). These models assume that the projections from retinal ganglion cells to cortical neurons pass through dLGN without modification. The emergence of orientation selectivity from random inputs is due to distance-dependent connectivity between them, and orientation columns naturally emerge in this scenario. In rodents, the orientation preferences of V1 neurons, however, do not seem to be neatly organized in patches and smooth maps (***Ohki et al., 2005***), and the salt-and-pepper distribution of preferences does not suggest spatial models to make a strong contribution.

It has been previously suggested (***Pattadkal et al., 2018***) that orientation selectivity could emerge from random connectivity, without a dedicated alignment of the sensors (***Hubel and Wiesel, 1962***), and in absence of orientation maps (***Ohki et al., 2005***). The explanation offered by Pattadkal and colleagues was that orientation-selective responses of cortical neurons could be the result of randomly emerging ON and OFF subfields of thalamic inputs. The weak and random orientation bias in the thalamic input would then be amplified by the excitatory-inhibitory cortical network. Under these conditions, it was found that the OSI was robust with regard to the number of thalamocortical projections and spatial frequency of the stimulus (apart from very low frequencies), while the PO itself depended strongly on spatial frequency. The exact input-output transformation was not considered. In contrast, our simulation results concluded that the orientation preference indeed depends on these parameters if a wider range of values is considered for them.

Adopting the same general idea in our new work, we have come up with a detailed explanation of the phenomenon by emphasizing other aspects of the computations performed by the thalamo-cortical circuit. In our model, we also assumed projections from thalamus to cortex with no particular *a priori* structure. Each individual cortical neuron thus extracts a different random sample of the visual field. If stimulated with a moving grating, the resulting compound input had a temporal modulation entrained by the grating, with a phase resulting from the interference of many oscillatory inputs of different phases. The amplitude of these resulting oscillations was tuned to orientation, in full agreement with experimental findings (***Lien and Scanziani, 2013***). Despite the same findings of the cortical OS dependence on the RF structures of thalamic inputs, our analysis revealed that the F1-to-F0 transformation, which is the key input-output transfer mechanism in our model, is naturally mediated by the nonlinear transfer performed by individual spiking neurons (see Fig 9G). Combining these two effects, our theory shows explicitly that the tuning of cortical neurons depends on the thalamo-cortical convergence number and on the spatial frequency of the stimulus (see Fig 11 and Fig 12). Compared to previous models, therefore, our model did not only exhibit reliable tuning in numerical simulations, but it also explained the detailed neuronal mechanisms underlying the emergence of contrast-invariant tuning curves in V1 neurons.

As the mechanisms described in our study are very general, they might also account for the emergence of feature selectivity in other sensory modalities, provided the information is conveyed in the amplitude of periodic signals. This might particularly apply to the whisker system in rodents, or the auditory system in all mammals.

## Methods and Materials

### Description of the model system

#### Network model

The basic model network used in this work is composed of two parts: the thalamic (dLGN) feedforward projection and the cortical (V1) recurrent network (***Figure 4***). The layout of our V1 network is identical to the one introduced by ***Brunel (2000)***. It consists of *N* = 12 500 leaky integrate-and-fire neurons, of which *a* = 80% are excitatory and 1 − *a* = 20% are inhibitory. The recurrent connectivity ϵ = 10% is uniform throughout the network (***Braitenberg and Schüz, 1998***). As a result, each neuron receives exactly 1 000 excitatory and 250 inhibitory inputs from within the same network, drawn randomly and independently. Self-connections are excluded. The amplitudes of excitatory recurrent synapses are *J*_E_ = 0.2 mV. Inhibitory couplings are set to be *g* = 8 times stronger than excitatory ones. As a consequence, the amplitudes of inhibitory synapses are *J*_I_ = −*gJ*_E_ = −1.6 mV. This results in an inhibition-dominated recurrent network.

Besides recurrent input, V1 neurons receive additional feedforward input from three sources: constant background, thalamic excitation and feedforward inhibition. Background inputs represent projections from any other brain areas except visual thalamus. In our model, they are identical for all recurrent neurons and keep the recurrent neural activity going in absence of visual stimulation. They are rendered as a stationary Poisson process with constant rate *v*_bg_. The synaptic weights are *J*_bg_ = 0.1 mV for all simulations. The second input source is the visual thalamus. In the primary visual cortex, the visual information is mainly conveyed by the dorsal lateral geniculate nucleus (dLGN) through thalamocortical projections. A recurrent V1 neuron receives input from exactly 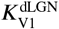 neurons in dLGN. As a result, the emerging receptive fields of recurrent V1 neurons are of similar size, but vary slightly from neuron to neuron. Receptive fields are roughly circular, but non-uniform, reflecting the random positions of dLGN inputs. Experiments *in vitro* reported that thalamocortical synapses are several times stronger than intracortical synapses (***Gil et al., 1999***; ***Richardson et al., 2009***). Here, we assume that the efficacy of direct thalamocortical projections is 10 times larger than recurrent excitatory connections,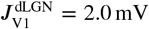. The third source of input are feedforward inhibitory projections (FFI) from other cortical neurons. FFI neurons represent a specific type of inhibitory interneurons, which selectively target recurrent V1 neurons. They receive input from 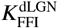 neurons in dLGN, each with a synaptic efficacy of 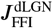. Thalamic afferents are the only driver of FFI neurons in our model, so their activity is fully determined by its thalamic inputs. Finally, 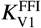 FFI neurons project onto each V1 neuron, each with synaptic weight 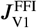. In thecortical circuit just described, the connections between FFI neurons and recurrent V1 neurons are established randomly and independently.

#### Neurons and receptive fields

Neurons in the lateral geniculate nucleus (dLGN) have circular antagonistic center-surround receptive fields, either ON-center/OFF-surround or OFF-center/ON-surround. ON-center cells respond strongest when the center of their receptive fields is exposed to light, and they are inhibited when the surround is illuminated. OFF-center cells respond in exactly the opposite way. The center and surround sub-regions of the receptive field are described by two-dimensional normalized Gaussian functions of different widths *σ*_+_ and *σ*_−_, respectively. For ON-center cells, we have *σ*_+_ < *σ*_−_ and *σ*_+_ > *σ*_−_ for OFF-center cells. The receptive field is simply represented by the difference of these two Gaussians (DoG) and has the form

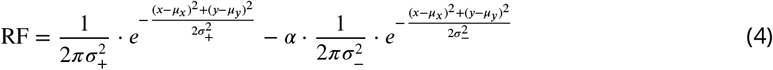

where (*μ*_*x*_, *μ*_*y*_) denotes the position of the receptive field center of dLGN neurons. The scaling factor *α* describes the relative weight of integrated subfields. In cat and monkey, the value is reported to be approx. 0.85 for retinal ganglion cells and LGN neurons (***Tadmor and Tolhurst, 2000***). In mouse superior colliculus neurons, the factor is approx. 1.07 (***Wang et al., 2010***). In our simulation, the scaling factor *α* is set to 1.0, indicating that the center and surround subfields are equally weighted. Note that our conclusions are not affected by any specific choice of this number. ***Figure 14*** shows an example of the receptive field of an ON center cell. In our model, an equal number of ON center and OFF center cells are distributed randomly in the visual field.

**Figure 14.**
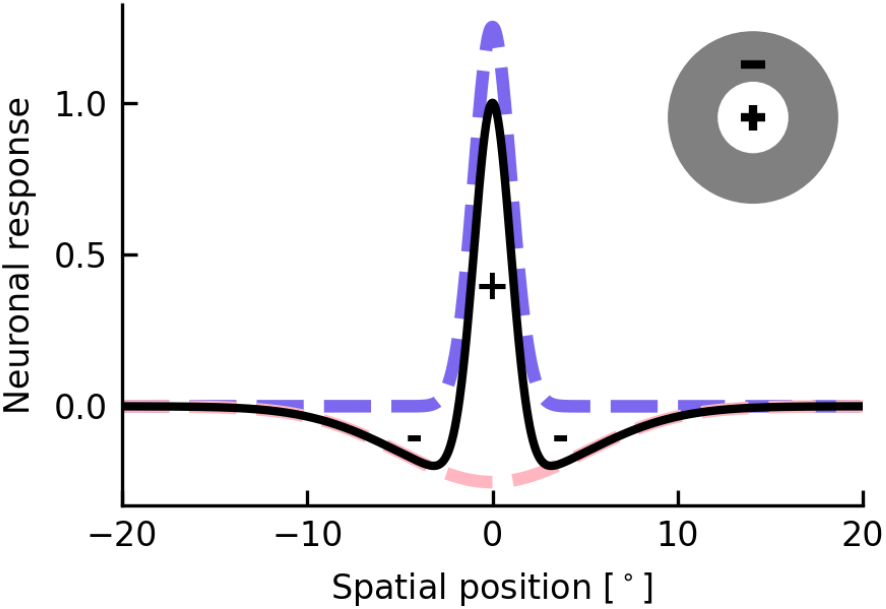
Isotropic receptive field of dLGN neurons. The two-dimensional receptive field of a single dLGN neuron is conceived as difference of Gaussians (DOG). Shown is the neuronal response (solid line) of an ON-center/OFF-surround cell (inset) to small spots of light at the position indicated. Dashed lines represent the responses for a separate stimulation of either the center or the surround, respectively.

In order to investigate orientation selectivity in line with experiments, we use moving oriented gratings with luminance changing sinusoidally both in space and time. Each of these visual stimuli has an orientation *θ*, a temporal frequency *f* and a spatial frequency *λ*. The movement direction of the grating is always orthogonal to its orientation. The light intensity of the stimulus at position (*x, y*) at time *t* is given by

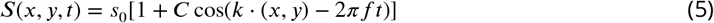

where *s*_0_ is the mean luminance of the stimulus, *C* is the contrast of the grating, and *k* = 2*πλ*(cos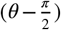, sin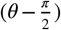) is the wave vector. In some works however, in contrast to our definition, *θ* denotes the direction of movement which is perpendicular to the stripes of the grating (***Pattadkal et al., 2018***; ***Kondo et al., 2016***). In this case, the wave vector becomes *k* = 2*πλ*(cos(*θ*), sin(*θ*)). The resulting firing rate of dLGN neuron *i* at position (*x*_*i*_, *y*_*i*_) in response to the stimulus grating can then be calculated as

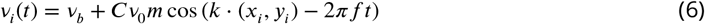

where 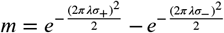 and *v*_0_ is the rate of a dLGN neuron in response to the mean stimulus luminance *s*_0_. *m*_max_ is the maximum value of *m* for a given set of (*σ*_+_, *σ*_−_) when varying the spatial frequency *λ*. The baseline firing rate is given by *v*_*b*_ = *m*_max_*v*_0_ such that the firing rate of a single dLGN neuron will always be non-negative.

The dLGN neurons are modeled as Poisson neurons, i.e. spikes are generated randomly and independently with firing rate *v*_*i*_(*t*) at each point in time. A Dirac delta-function 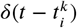 is used to represent spike *k* generated by neuron *i*. The spike train of dLGN neuron *i* is the sum of all spikes it generates 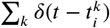.

Cortical neurons, in contrast, are conceived as leaky integrate-and-fire (LIF) neurons. The subthreshold time evolution of the membrane potential *V*_*i*_(*t*) of neuron *i* is determined by

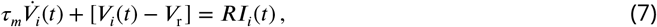

where *τ*_*m*_ is the membrane time constant and *R* is the leak resistance. The current *I*_*i*_(*t*) represents the total input to neuron *i*. A spike is elicited when the membrane potential reaches the threshold *V*_th_, after which *V*_*i*_(*t*) is reset to its resting potential *V*_r_. It remains at the resting potential for a short refractory period *t*_ref_. During this absolute refractory period, no spike will be generated.

### Mathematical implementation of the model

#### Network of spiking neurons

In order to study the orientation preference of recurrent neurons in the network described above, we set up a spiking neuronal network. In this network, cortical neurons receive presynaptic spike inputs, resulting in transient changes of the postsynaptic membrane potential. Excitatory and inhibitory recurrent neurons, as well as FFI neurons, are conceived as leaky integrate-and-fire neurons. The time evolution of the membrane potential *V*_*i*_(*t*) is described by a differential equation Eq 7, separately for each neuron *i*. Thereby, the total input current *I*_*i*_(*t*) is the superposition of all inputs

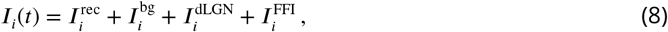

accounting for their respective synaptic strengths.

The background inputs 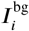 are all identical and target recurrent V1 neurons. They are modeled as a Poisson process with mean firing rate *v*_bg_. The corresponding background input current is given by

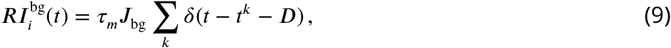

where *t*^*k*^ is the emission time of *k*-th spike of the input and *J*_*bg*_ is the amplitude of postsynaptic potential.

In all other pathways, convergent projections need to be accounted for. Therefore, the input current of each component is given by the sum of spike input from all presynaptic neurons, indexed by *j*

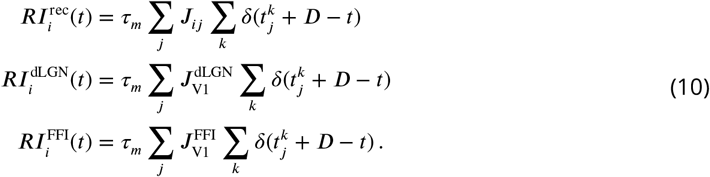

Note that this entails ϵ*N*, 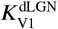 and 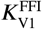 non-zero contributions to the respective sum, respectively. Although the convergence numbers are the same for all recurrent V1 neurons in each pathway, their presynaptic neurons are different. As the FFI neurons are also modeled as LIF neurons, the same method was be applied to calculate their input currents and then extract the respective spike trains.

Combining all the inputs above, the V1 neurons in the network respond to a visual stimulus in terms of spike trains. All numerical simulations of this model were performed in the neural simulation tool NEST (***Gewaltig and Diesmann, 2007***; ***Fardet et al., 2020***). All the parameters used in numerical simulations are shown in ***Table 1***.

**Table 1.** Table of parameters.

| **Network connectivity** |  |  |
| --- | --- | --- |
| Number of V1 neurons | $N$ | 12 500 |
| Recurrent connection probability | $\epsilon$ | 10% |
| Number of dLGN neurons | $N_{\text{dLGN}}$ | 3 071 |
| Number of FFi neurons | $N_{\text{FFi}}$ | 1 500 |
| Projection number of dLGN→V1 | $K_{\text{V1}}^{\text{dLGN}}$ | 100 |
| Projection number of dLGN→FFi | $K_{\text{FFi}}^{\text{dLGN}}$ | 8 |
| Projection number of FFi→V1 | $K_{\text{V1}}^{\text{FFi}}$ | 320 |
| **Neuron model** |  |  |
| Membrane time constant | $\tau_{\text{m}}$ | 20 ms |
| Refractory period | $\tau_{\text{ref}}$ | 2 ms |
| Membrane resistance | $R$ | 40 MΩ |
| Membrane capacitance | $C_{\text{m}}$ | 250 pF |
| Resting potential | $V_{\text{r}}$ | 0 mV |
| Threshold voltage of V1 neurons | $V_{\text{th}}$ | 20 mV |
| Threshold voltage of FFi neurons | $V_{\text{th}}^{\text{FFi}}$ | 20 mV |
| **Synaptic model** |  |  |
| Synaptic delay | $D$ | 1.5 ms |
| Recurrent exc. synaptic efficacy | $J_{\text{E}}$ | 0.2 mV |
| Inhibition dominance ratio | $g$ | 8 |
| Recurrent inh. synaptic efficacy | $J_{\text{I}}$ | $-gJ_{\text{E}}$ |
| Background input strength | $J_{\text{bg}}$ | 0.1 mV |
| Synaptic strength of dLGN→V1 | $J_{\text{V1}}^{\text{dLGN}}$ | 2.0 mV |
| Synaptic strength of dLGN→FFi | $J_{\text{FFi}}^{\text{dLGN}}$ | 2.0 mV |
| Synaptic strength of FFi→V1 | $J_{\text{V1}}^{\text{FFi}}$ | -1.6 mV |
| **Simulation** |  |  |
| Background firing rate | $\nu_{\text{bg}}$ | 8 350 Hz |
| Mean luminosity of grating | $s_0$ | |
| dLGN rate response to $s_0$ | $\nu_0$ | 100 Hz |
| Stimulus orientation | $\theta$ | 0°, 15°, ... , 165° |
| Spatial frequency | $\lambda$ | 0.04, 0.08, ... cpd |
| Temporal frequency | $f$ | 3 Hz |
| Simulation time | $T$ | 6 s |
| Contrast | $C$ | 0, 0.1, 0.3, 0.5, 0.8, 1 |

#### Analytical firing rate model

Although spiking neurons (here, LIF) provide a more biologically realistic model, the numerical effort to study input-output transfer functions via simulations is quite high. To reduce the effort, and to provide additional mathematical insight, we employed analytical firing rate models that generalize the well-known diffusion approximation to certain time-dependent inputs. First, we devised a stationary rate model (SRM) to estimate the output. Assuming that the input to the neuron changes slowly, allowing that it is in equilibrium in every moment, we can just use the known steady-state solution, moment by moment. However, this approach has limitations. As in the setting considered here V1 neurons receive oscillatory inputs from the thalamus, a dynamic rate model (DRM) appeared to be more appropriate. As the membrane acts as a lowpass with frequency-dependent attenuation, we assume that the input amplitude depends on the temporal frequency *f* according to 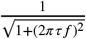, where *τ* is the time constant of the membrane. On this basis, the DRM provides good estimates of the output firing rate for a wider range of temporal frequencies.

The nonlinear firing rate model used here is based on the diffusion approximation to single neurons, see e.g. ***Brunel (2000)***; ***Siegert (1951)***; ***Ricciardi (1977)***; ***Amit and Tsodyks (1991)***. In this setting, the total synaptic input current to a neuron *i* is replaced by a Gaussian White Noise of mean *μ*_*i*_ and amplitude *σ*_*i*_, which drives the neuron in an equivalent way

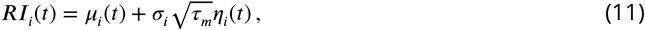

As described above, the total input to a single recurrent V1 neuron is composed of the recurrent and feedforward inputs from four sources. Assuming their statistical independence, this yields

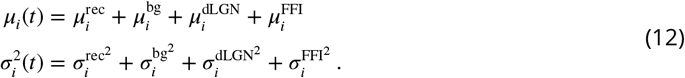

Mean and variance of each source is related to the respective presynaptic firing rates *v* and synaptic strength *J*. When a sinusoidal grating moves over the visual field, the firing rates of all individual dLGN neurons change sinusoidally over time (see Eq 6). As the rate of background input is fixed for all recurrent V1 neurons, the mean and variance of the background input is constant over time. The rates of FFI neurons are determined by their thalamic inputs. The mean and variance of each part is then given by

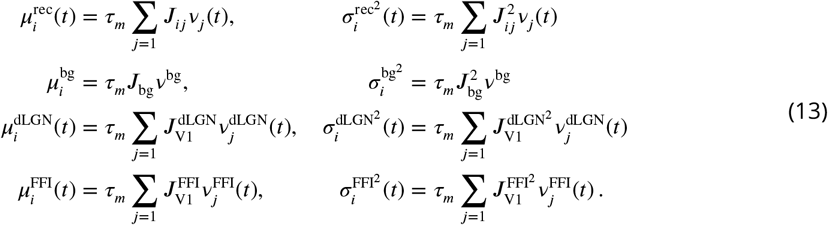

The steady-state firing rate of all recurrent V1 neurons are given by a transfer function *F*_*i*_ ***Siegert (1951)***

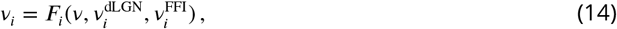

where *F*_*i*_ is defined by

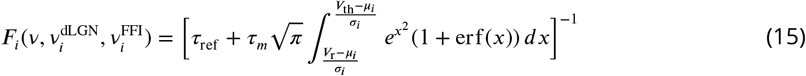

where erf is the error function. The self-consistent solutions of these nonlinear equations will return the estimation of the firing rates of individual recurrent neurons.

The firing rates of FFI neurons are easier to obtain, as there are no recurrent connections and we do not need to solve the equations self-consistently. As each FFI neuron *i* only receives inputs from dLGN neurons, its input mean and variance is given by

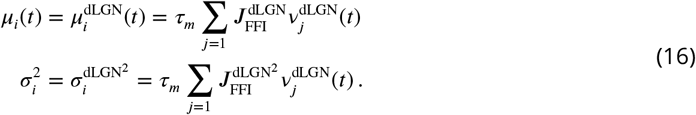

The steady state firing rate of FFI neuron *i* can be calculated from the transfer function 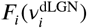 by solving Eq 15 with their respective inputs.

### Visual Stimulus

#### Drifting Grating

Sinusoidal drifting gratings were used as stimuli to extract the orientation tuning of neurons. They covered the entire visual field. The gratings were presented at 12 different orientations, evenly covering the range between 0° and 180° in discrete steps of 15°. The movement direction of the grating was always orthogonal to the orientation of the grid lines. Each stimulus lasted for 6 s.

#### Receptive field

In order to better compare our network simulations to experimental results, two different stimulation protocols were adopted to measure the receptive fields. In the Results, the protocol is similar to the experiments in ***Lien and Scanziani (2013)***. Each stimulus consists of a light (maximum luminance) or dark (minimum luminance) square on top of a gray (mean luminance) background. The width of each square is 5°, and it is randomly placed at one of 35 × 35 locations to cover the entire 175° × 175° stimulus field. Note that the actual width of the visual field is only 134°. The stimulus field for calculating the receptive fields of neurons is extended to 175° to avoid boundary effects. Each stimulus is presented for 20 s, its location and luminance (light or dark) are random. Each location of the grid is eventually stimulated with light and dark squares. The total stimulation time is 49 000 s.

The receptive fields of cortical recurrent neurons and their respective thalamic inputs are also mapped using locally sparse noise (see Appendix). For each stimulus image, an equal number of light and dark spots are placed randomly in the visual field on a gray background. Approximately 20% of the visual field is covered by spots. Again, in order to eliminate the boundary effects of dLGN neurons at the border of the visual field, the stimulus image is extended by gray background. The diameter of the spots is 4° and the resolution of the grid for positioning is 0.2°. In total, 20 000 stimulus frames are used during a simulation, and each frame is presented for 33 ms.

### Data analysis

#### Orientation selectivity

To quantify the orientation selectivity of single cortical neurons, the preferred orientation (PO) and the orientation selectivity index (OSI) are calculated for each neuron. This information can be extracted from its respective tuning curve, *v*(*θ*), representing the mean firing rate of a neuron for stimulus orientation *θ*. The method used here is to first compute the orientation selectivity vector (here represented as a complex number) from circular statistics (***Batschelet et al., 1981***; ***Piscopo et al., 2013***)

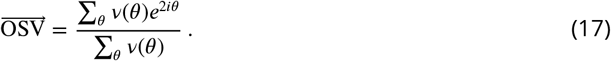

The PO is then extracted as the phase (angle) of the OSV

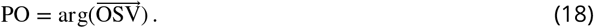

In contrast, the OSI is extracted as the magnitude (length) of the OSV

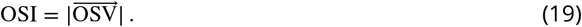

The OSI is often used to describe the strength of orientation selectivity. A neuron with high orientation selectivity, which only responds to one stimulus orientation and keeps silent for other orientations, returns OSI = 1. For an unselective neuron responding to all orientations equally, we have OSI = 0.

In some experimental literature, an alternative measure of orientation selectivity is used. It is calculated by OSI^*^ = (*v*_pref_ −*v*_orth_)/(*v*_pref_ +*v*_orth_), where *v*_pref_ is the firing rate at the preferred orientation and *v*_orth_ is the firing rate at its orthogonal orientation. In previous theoretical work, it has been pointed out that, for a perfect cosine tuning curve, OSI^*^ is twice as large as the OSI (***Sadeh et al., 2014***).

#### Preferred orientation of gratings

In order to evaluate the comparison of the orientation preference across different conditions (e.g. spatial frequency of the stimulus), we use the circular correlation (CC) of PO (***Pattadkal et al., 2018***). If PO_*i*_ is the preferred orientation of neuron *i* at one spatial frequency, *θ*_*ij*_ = PO_*i*_ − PO_*j*_ is the difference of PO of neuron *i* and *j* at this spatial frequency, and 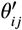 is the difference of neuron *i* and *j* at another spatial frequency. The circular correlation between PO at different spatial frequencies is extracted by

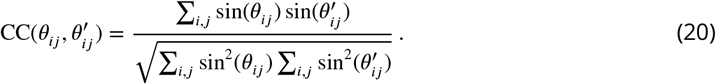

The value of CC ranges from −1 to 1. The preferred orientations at different spatial frequencies are perfectly linear correlated when CC = 1, and CC = 0 means no correlation between them.

#### Receptive fields

The raw receptive fields are estimated by reverse correlation. This is a commonly used method to reconstruct the receptive fields by averaging all the frames, each of them weighted by the neuronal response it evokes. By following the method described in ***Lien and Scanziani (2013)***, we extract the ON and OFF subfields of V1 neurons and their thalamic inputs and then predict the PO of the receptive fields (RF_Pref_). As shown in ***Figure 8***, the RF_Pref_ is orthogonal to the axis connecting the peaks of the ON and OFF subfields, respectively. The similarity between the RF of V1 neuron and its compound thalamic input is calculated as the correlation coefficient between them. The raw receptive fields of dLGN neurons, V1 neurons and their thalamic inputs stimulated by sparse noise are shown in Appendix. Note that no extra smoothing was applied to these figures of receptive fields.

## Acknowledgments

This work was partially funded by the joint training program SMARTSTART, launched by the Bernstein Network Computational Neuroscience and the Volkswagen Foundation. Additional support was obtained from the European Union’s Seventh Framework Programme (FP7/2007-2013) under Grant Agreement 600925 (NeuroSeeker) and from the DFG (grant EXC 1086). The HPC facilities used for this work are funded by the state of Baden-Württemberg through bwHPC and DFG grant INST 39/963-1 FUGG. In addition, our work was promoted by the German Academic Exchange Service (DAAD) and the Carl Zeiss Stiftung. The funders had no role in study design, data collection and analysis, decision to publish, or preparation of the manuscript.

We thank Uwe Grauer from the Bernstein Center Freiburg as well as Bernd Wiebelt and Michael Janczyk from the Freiburg University Computing Center for their assistance with HPC applications and the developers of the neural simulation tool NEST. We also thank Philippe Coulon and Nebojša Gašparović for their valuable advises.

## Appendix

### Appendix 1

**Figure S1.**
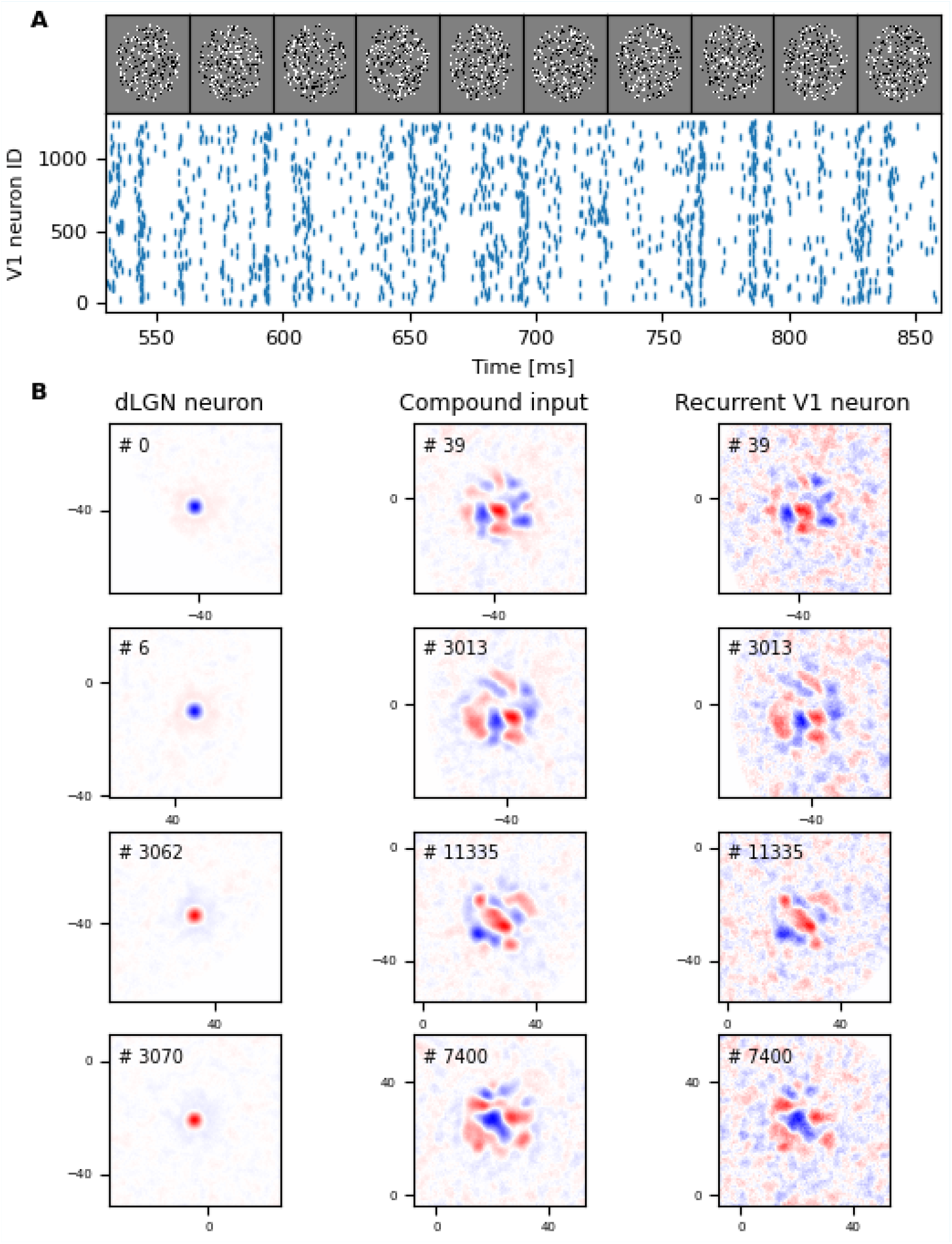
The neuronal responses and receptive fields stimulated by sparse noise. **A** Top row shows sample frames of a sparse spot stimulus. Each frame is presented for 33 ms. The bottom raster plots are the spikes of V1 neurons (10% of the whole population) that were elicited by sparse spot stimulation. **B** Some examples of receptive fields stimulated by sparse noise shown in **A**. The receptive fields of four example dLGN cells (2 ON center and 2 OFF center cells) are shown in the first column. The right and middle column depict the RFs of four V1 neurons and their thalamic inputs, respectively. The example neurons are exactly the same as in ***Figure 8***. Blue and red colors indicate ON and OFF subfields.

